# Time course of homeostatic structural plasticity in response to optogenetic stimulation in mouse anterior cingulate cortex

**DOI:** 10.1101/2020.09.16.297606

**Authors:** Han Lu, Júlia V. Gallinaro, Claus Normann, Stefan Rotter, Ipek Yalcin

**Author notes:** Shared corresponding authors: Stefan Rotter, Bernstein Center Freiburg & Faculty of Biology, University of Freiburg, 79104 Freiburg, Germany; Hansastraße 9a, 79104 Freiburg, Germany; Ipek Yalcin, Centre National de la Recherche Scientifique, Université de Strasbourg, Institut des Neurosciences Cellulaires et Intégratives UPR3212, 67000 Strasbourg, France; 8 Allée du Général Rouvillois, 67000 Strasbourg, France.

## Abstract

Plasticity is the mechanistic basis of development, aging, learning and memory, both in healthy and pathological brains. Structural plasticity is rarely accounted for in computational network models, due to a lack of insight into the underlying neuronal mechanisms and processes. Little is known about how the rewiring of networks is dynamically regulated. To inform such models, we characterized the time course of neural activity, the expression of synaptic proteins, and neural morphology employing an *in vivo* optogenetic mouse model. We stimulated pyramidal neurons in the anterior cingulate cortex of mice and harvested their brains at 1.5 h, 24 h, and 48 h after stimulation. Stimulus-induced cortical hyperactivity persisted up to 1.5 h and decayed to baseline after 24 h, indicated by c-Fos expression. The synaptic proteins VGLUT1 and PSD-95, in contrast, were upregulated at 24 h and downregulated at 48 h, respectively. Spine density and spine head volume were also increased at 24 h and decreased at 48 h. This specific sequence of events reflects a continuous joint evolution of activity and connectivity that is characteristic of the model of homeostatic structural plasticity. Our computer simulations thus corroborate the observed empirical evidence from our animal experiments.

## Introduction

Neural circuits in the mammalian brain are highly plastic (Holtmaat and Svoboda 2009). Synaptic plasticity comes in two different flavors. Functional plasticity means that chemical synapses change their strength by modifying signal transmission based on neurotransmitters and receptors (Bear and Malenka 1994; Malenka and Bear 2004). Structural plasticity, in contrast, refers to a variety of changes including the branching of dendrites, the geometry of dendritic spines, and number of dendritic spines and axonal boutons, and the connectivity between specific pairs of neurons (Trachtenberg et al. 2002; Caroni et al. 2012; Pfeiffer et al. 2018). Both forms of plasticity are underlying network assembly during development, use-dependent adaptation and learning in the adolescent and adult brain, but also network decay during aging and disease (Lamprecht and LeDoux 2004). Memory depends on plasticity. For instance, fear conditioning has been shown to increase both the synaptic strength and connection probability among a subgroup of granule cells in the dentate gyrus (Ryan et al. 2015). The resulting memory engram encodes a distinct episodic memory (Liu et al. 2012; Ramirez et al. 2013). Plasticity caused by injury, such as synaptic potentiation and network remodeling triggered by stroke or brain lesion, is likely to involve both activity perturbation and neuroinflammation (Keck et al. 2008; Murphy and Corbett 2009). In brain diseases, pathological plasticity may affect several brain regions. Acute and chronic stress, for instance, has been shown to induce different functional and structural alterations in the hippocampus, anterior cingulate cortex (ACC), amygdala (Lucassen et al. 2014), and elsewhere. Given this wealth of phenomena, the question arises how functional and structural plasticity is regulated.

The rules underlying experience-dependent plasticity need to be investigated further. Experiments in different brain regions with different plasticity-inducing paradigms have given rise to a host of different phenomena (Bear and Malenka 1994; Malenka and Bear 2004; Keck et al. 2017; Gainey and Feldman 2017). Correlation-based Hebbian plasticity, summarized as “neurons that fire together wire together” (Hebb 1949), was proposed to account for homosynaptic strengthening observed in animals minutes to hours after artificial high-frequency stimulation (Lowel and Singer 1992). Despite its great potential in explaining learning and memory, Hebbian plasticity in computational network models was shown to increase the risk of excessive excitation or silencing, respectively (Sejnowski 1977; Miller and MacKay 1994). The same lack of network-level stability is implied by spike timing dependent plasticity (STDP), which implements either homosynaptic strengthening or weakening based on the relative timing of presynaptic and postsynaptic spikes (Markram et al. 1997; Bi and Poo 1998). The discovery of heterosynaptic plasticity and synaptic scaling, however, hinted that the modulation of a synapse may also depend on its neighbors (Lynch et al. 1977; Chater and Goda 2020) and the activity of the postsynaptic neuron (Turrigiano 2012). Chronic *in vivo* recordings have indeed revealed a robust cell-by-cell firing rate homeostasis across days and weeks (Hengen et al. 2016; Ma et al. 2019; Pacheco et al. 2019). New models of homeostatic plasticity (Turrigiano 2012, 2017), possibly in combination with Hebbian plasticity rules, are now being evaluated for their ability to solve the aforementioned network stability issues. Preliminary conclusions posit that the time scales of homeostatic control should be much faster than those observed in experiments (Zenke and Gerstner 2017). Rarely, however, was a possible role of structural plasticity explored in these theoretical studies.

Structural and functional plasticity do not represent independent processes. Changes in spine number and individual spine head volumes were observed after synaptic potentiation or depression *in vitro* (Engert and Bonhoeffer 1999; Matsuzaki et al. 2004; Zhou et al. 2004). Moreover, both spine density and synaptic strength were shown to compensate for input loss caused by entorhinal denervation in organotypic slice culture (Vlachos et al. 2012b; Lenz et al. 2019). In parallel, theoreticians began to reflect over possible functional aspects of structural plasticity in a network (Fauth and Tetzlaff 2016). The homeostatic structural plasticity (HSP) model seems particularly promising in reconciling robust development and associative learning (Butz et al. 2009; Van Ooyen 2011; Butz and van Ooyen 2013; Gallinaro and Rotter 2018; Lu et al. 2019; Gallinaro et al. 2020). Still, the empirical data justifying such activity-dependent structural plasticity models are sparse. Most studies report changes in spine density after massive manipulation of activity, or in brain diseases, see review by Chidambaram et al. (2019). Unfortunately, time-resolved neural activity and connectivity was not included in any of them. Yusifov et al. (2021) revealed the elaborate temporal dynamics of spine density during monocular deprivation but only apical dendrites were monitored. The time course of structural changes while the neuronal activity recovers, however, is of great importance to disambiguate structural plasticity models.

The optogenetic stimulation paradigm employed in the present study evolved from our previous studies. We activated the pyramidal neurons in mice ACC for four consecutive days (30 minutes per day) until they developed depressive-like behaviors. Using the same optogenetic stimulation paradigm, we sampled mouse brains at 1.5 h, 24 h and 48 h after chronic stimulation. We stained and quantified the relative abundance of neuronal activity marker c-Fos, and general synaptic markers the vesicular glutamate transporter 1 (VGLUT1) and the postsynaptic density scaffold protein PSD-95 (De Gois et al. 2005; Ehrlich et al. 2007). As expected, optogenetic stimulation triggered depressive-like behaviors and hyperactivity in mice (Barthas et al. 2015). Hyperactivity of pyramidal neurons, evidenced by a robust c-Fos expression at 1.5 h, eventually diminished to baseline 24 h after the end of the stimulation, while VGLUT1 and PSD-95 showed strong delayed upregulation at 24 h and again downregulated after 48 h in the stimulated mice. Similar to the temporal expression profile of VGLUT1 and PSD-95, dendritic spine density and spine head volume of the stimulated mice were increased at 24 h and restored to the control levels, or even slightly below control at 48 h. Microglia and astrocytes are known to be involved in structural plasticity (Weinhard et al. 2018) and glutamate homeostasis (Schousboe and Waagepetersen 2005). Therefore, we also checked whether glial markers, GFAP and IBA1, were overexpressed throughout 48 h after stimulation. We argue that, compared to other candidate theories, the homeostatic structural plasticity model could explain the biphasic changes of synaptic proteins and dendritic morphology consistently at 24 h and 48 h after the chronic stimulation.

## Materials and Methods

### Animals

3*−*5 months old genetically modified mice expressing ChR2 and yellow fluorescent protein (Thy1-ChR2-YFP) in a subset of pyramidal neurons (MGI Cat# 3719993, RRID:MGI:3719993) as well as C57BL/6J male adult mice (IMSR Cat#JAX:000664, RRID:IMSR JAX:000664; Charles River, L’Arbresle, France) were used in the current study. All mice were kept in a reversed day-night cycle, with food and water provided *ad-libitum*. Mice were firstly group caged and then single housed after the optic fiber implantation. The Chronobiotron animal facilities are registered for animal experimentation (Agreement A67-2018-38), and protocols were approved by the local ethical committee of the University of Strasbourg (CREMEAS, *n^◦^* 02015021314412082).

### Animal experimental design

The animal experiments’ objective was to determine the time course of plastic phenomenon triggered by external stimulation. We adopted an established optogenetics mouse model from our laboratory, in which the pyramidal neurons in ACC (24a/24b) were activated for four consecutive days (details see Optogenetic stimulation section below). We studied the temporal dynamics with discrete time points, by harvesting the mouse brain tissue at 1.5 h, 24 h, or 48 h after the last stimulation. As shown in our previous studies (Barthas et al. 2015, 2017), sustained stimulation of the ACC induces depressive-like behavior in näıve mice. So in the current study, we used splash test and novelty-suppressed feeding (NSF) test to verify the behavioral effects of the optogenetic stimulation. In the 24 h-post groups, we conducted splash test on the fifth day, while in the 48 h-post groups, we performed both NSF test and splash test on the fifth and sixth day and sacrificed the mice afterwards. We later evaluated the temporal evolution of neural activity by quantifying the expression of c-Fos at the three aforementioned time points. To capture when and where synaptic alterations may occur, the expression of pre- and postsynaptic proteins were evaluated. Following the pattern shown by preliminary molecular screening, we examined if structural changes accompany molecular alterations by estimating the spine morphology of ACC pyramidal neurons harvested at 24 h and 48 h post-stimulation. To further confirm the involvement of glial cells, the expression of glial markers at 1.5 h, 24 h, and 48 h were respectively inspected. Before we performed all the experiments in Thy1-ChR2-YFP mice, we compared the efficacy of the transgenic approach with viral transfection. For the latter, we injected bilaterally AAV-CaMKII-ChR2 (H134R)-EYFP (Addgene plasmid #26969; http://n2t.net/addgene:26969; RRID: Addgene 26969) into the ACC (details see Virus injection section in the Supplementary Materials) of C57BL/6J mice. As we observed no differences in the c-Fos activity and behavioral outcomes in two approaches after optogenetic activation, we decided to perform all the experiments in transgenic mice to reduce the number of surgery that animals go through. All mice group information was summarized in Supplementary Table 1.

### Stereotactic surgery

Stereotactic surgery was conducted to inject virus and implant optic fiber into ACC. During the surgery, mice were deeply anesthetized with a mixture of zoletil (25 mg*/*kg tiletamine and 25 mg*/*kg zolazepam) and xylazine (10 mg*/*kg) (Centravet, Taden, France; i.p. injection) and locally anesthetized by bupivacaine (Mylan, The Netherlands; 0.5 mg*/*mL; subcutaneous injection, 1 mg*/*kg). The coordinates of the injection/implantation site are

+0.7 mm from bregma, lateral: *±*0.3 mm, dorsoventral: *−*1.5 mm from the skull (Barthas et al. 2015; Sellmeijer et al. 2018).

### Optic fiber implantation

We inserted 1.7 mm-long LED optic fiber (MFC 220/250-0.66 1.7mm RM3 FLT, Doric Lenses, Canada) unilaterally (left or right) in C57BL/6J mice two weeks after the virus injection or directly in näıve Thy1-ChR2-YFP mice. The fiber was inserted into ACC for 1.5 mm deep with reference to the skull. The metal end was fixed onto the skull by superglue and dental cement, and then the skin was stitched. For stimulation, we used blue light (460 nm wavelength) and the light intensity of optic fibers used in the current study ranged from 1.7 mV*/*mm^2^ to 6 mV*/*mm^2^.

### Optogenetic stimulation

After the optic fiber implantation, we individually housed the mice to avoid possible damage to the implant. After seven days of recovery, we started the optogenetic stimulation protocol on freely moving mice in their home cages. Optogenetic stimulation took place on four consecutive days. We applied repetitive pulses for a total duration of 30 minutes, in which the pulses were organized into multiple 10 seconds long pulse trains, each consisting of 8 seconds at 20 Hz with 40 ms pulse duration and 2 seconds without stimulation. We did not observe the effects of light on the behaviors in gene-matched wild type mice (Supplementary Figure 1-2). We used transgenic mice for all experiments and kept the light off for the sham groups. At the end of the stimulation, all mice were handled again and unplugged from the cable.

### Behavioral tests

We performed all the behavioral tests during the dark phase under red light. Splash test (Nollet et al. 2013) and novelty-suppressed feeding (NSF) test (Samuels and Hen 2011) were used to evaluate depressive-like behaviors. In the splash test, we sprayed 15% sucrose solution onto the coat of the mice and recorded the total grooming time for each mouse during the following 5 min. The NSF test was conducted on a different day of splash test and we removed the food pellets 24 h before testing. During the test, we put each mouse into an open field, where a food pellet was placed in the middle, and recorded the time delay necessary for each mouse to touch and eat the pellet (within 5 min).

### Verification of injection site and tissue harvesting

Mice were perfused with cold 4% paraformaldehyde (PFA) in 1*×* phosphate buffer (PB, 0.1 M, pH 7.4), under Euthasol Vet (intraperitoneal injection, 2 *µ*L*/*kg; TVM, UK) overdose anesthesia. The details of timing and pump speed can be found in the Supplementary Materials. Frontal sectioning of the brains (40 *µ*m-thick for immunohistochemical staining and 300 *µ*m-thick for microinjection) was performed on a vibratome (Leica-VT1000s, Rueil-Malmaison, France). The injection or implantation site of each perfused mouse was checked under the microscope.

### Immunohistochemical staining

We did fluorescent staining to examine the expression of c-Fos, VGLUT1, PSD-95, Neurogranin, and GFAP (see Supplementary Table 2 for the antibody concentrations). We used sections ranging from 1.42 mm to *−*0.23 mm away from Bregma, with a distance 160 *µ*m in between. The sections were firstly washed in 1*×* PBS (3 *×* 10 min) and then blocked at room temperature (RT) with 5% donkey serum in 0.3% PBS-T (1 h). Later the sections were incubated at 4 °C with corresponding primary antibody and 1% donkey serum in 0.3% PBS-T overnight. Sections were rinsed with 1*×* PBS (3 *×* 10 min) in the next morning, incubated with secondary antibody in 0.3% PBS-T at RT (2 h), and rinsed again with 1*×* PBS (3 *×* 10 min). Sections were mounted on gelatin-coated slides, air-dried, and coverslipped with Vectashield H-1000 (Vector Laboratories, Germany).

We stained IBA1 with 3, 3*^t^*-Diaminobenzidine (DAB, Sigma, US). Sections were selected, washed, blocked, and treated with primary and secondary antibodies as described above. Then the sections were rinsed with 1*×* PBS (3 *×* 10 min) and incubated with avidin–biotin–peroxidase complex (ABC Elite, Vector Laboratories, Germany; 0.2% A and 0.2% B in 1*×* PBS) at RT (1.5 h). Later the sections were rinsed with 0.05 M Tris-HCl buffer (TB; pH 7.5; 3 *×* 10 min). Peroxidase revelation was achieved by incubation shortly (20 s) with a mixture of 0.025% DAB and 0.0006% H_2_O_2_ in 0.05 M TB. Sections were carefully rinsed with TB (2 *×* 10 min) and 1*×* PBS (2 *×* 10 min) to cease the reaction. All sections were mounted and air-dried, then dehydrated in graded alcohol baths (1 *×* 5 min in 70%, 1 *×* 5 min in 90%, and 2 *×* 5 min in 100%), cleared in Roti-Histol (Carl Roth, Germany), and coverslipped with Eukitt.

### Microinjection

We used microinjection and confocal microscope (Dumitriu et al. 2011) to visualize and quantify the neural morphology at 24 h and 48 h post-stimulation. The sections were selected within the distance of approximately *±*0.4 mm anterior-posterior (AP) away from the optic fiber. The injection was done only into the pyramidal neurons from layer 2 *−* 3 of ACC (24a/24b) from both hemispheres. The injection pipettes were pulled from glass capillaries with filament, with a final resistance around 150 MΩ. We filled the pipette with red fluorescent dye solution Alexa 568 hydrazide (#A10441, Thermo Fisher, USA) in filtered 1*×* PBS (1 : 40). We performed microinjection under the microscope of a patch-clamp set-up. During injection, we penetrated the pipette tip into the soma and switched on the current to *−*20 pA to drive the dye diffusion for 20 min. Later we switched off the current but left the pipette tip inside the soma for another 5 min to fill the dendrite and spines. All the sections were retrieved and covered with Vectashild H-1000 (Vector Laboratories, Germany) for confocal microscope imaging. We checked all the injected neurons for YFP signal; only neurons with YFP signal were identified as pyramidal neurons and selected for further analysis.

### Microscope imaging

To quantify the expression of c-Fos, VGLUT1, PSD-95, neurogranin, and GFAP in the ACC, we imaged epifluorescent signals of stained sections with Morpho Strider on Zeiss Imager2 (Carl Zeiss, Germany) with 2.5*×* objective. To achieve better resolution of representative images, we also imaged the sections at the middle focal plane with a confocal microscope Leica SP8 (Leica Microsystems, Germany; software Leica SP8 LAXS 3.5.6) with objectives HCX PL Fluotar 5 *× /*0.15, 20*×*, and HC PL APO CS2 63 *× /*1.40. The bright-field images of DAB-stained IBA1 were acquired with a NikonEclipse E600 microscope with 4*×* and 40*×* objectives (MBF Bioscience, USA; software Neurolucida 2019).

To analyze the morphological features, we took z-stacked images of microinjected neurons (with step size 0.2 *µ*m-0.3 *µ*m sampled by the software Leica SP8 LAXS 3.5.6) with confocal microscope Leica SP8 (Leica Microsystems, Germany). The entire neuron structure was imaged with the objective HC PL APO CS2 63 *× /*1.40. If not stated otherwise, we used a pulsed laser for excitation (White Light; 488 nm). Segments of apical and basal dendrites of each neuron were imaged with the objective HC PL APO CS2 63 *× /*1.40 as well. Second and third level dendrite segments with little overlap and clear background were selected.

### 3D reconstruction and analysis of dendritic morphology

Firstly, after imaging, we deconvolved our confocal z-stack images with Huygens Professional 19.04 (Scientific Volume Imaging, The Netherlands) to restore the object from the acquired image through knowledge of the point spread function (PSF) and noise. 3D reconstruction and morphological analysis were later performed on the deconvoluted images.

For each pyramidal neuron, we reconstructed the soma and its dendritic tree with Imaris 9.5.1 (ImarisXT, Bitplane AG, Switzerland). Based on the reconstructed data, the dendritic tree structure was represented by Sholl intersections (Sholl 1953) at different radiuses. The order of each dendrite segment and its corresponding length and average diameter were also estimated. We further used Fiji (ImageJ, Fiji) to measure the soma size of each neuron on its z-projected image.

We reconstructed the dendritic shafts and spines with Imaris 9.5.1 again for selected dendrite segments at high resolution. We also classified the spine classes (filopodia, long-thin, stubby, and mushroom) based on their morphological features with the Imaris Spines Classifier package. The criteria of spine classification were summarized in Supplementary Table 3. We harvested the overall spine density of each segment and the spine density of each spine class based on the reconstructed data. The spine head volume of individual spines was also estimated.

### Quantifying immunohistochemical staining images

Visually inspection showed the expression of marker proteins was not homogeneously distributed in ACC but constrained to the vicinity around the optic fiber. To reflect such a pattern, we systematically analyzed the expression of c-Fos, VGLUT1, PSD-95, Neurogranin, GFAP, and IBA1 in both hemispheres at different distances to the optic fiber at 1.5 h, 24 h, and 48 h post-stimulation.

We firstly organised the corresponding epifluorescent images (obtained under 2.5*×* objective) or bright-field images (obtained under 4*×* objective) of each marker for each mouse in sequential order. The Bregma level of each section was identified in reference to the Mouse Brain Atlas (Franklin et al. 2008). Later we checked the implantation site for each section. Sections with a clear trace of implantation were marked as *distance zero*. Sections at a more anterior position than the distance zero were labeled with a negative sign (*−*), while posterior sections were labeled with a positive sign (+). In the end all sections were classified into five distance groups and their average distances were noted as *−*0.4 mm, *−*0.2 mm, 0 mm, +0.2 mm, and +0.4 mm. Both hemispheres were also carefully identified as the ipsi- or contralateral side in reference to the implantation site.

To quantify the signal intensity of markers on each section, we created two same-sized masks (700 *µ*m *×* 700 *µ*m) on both hemispheres with Fiji. For c-Fos and IBA1, we counted signal positive cell numbers within each section’s masks, while for VGLUT1, PSD-95, neurogranin, and GFAP, we quantified the fluorescent intensity within the masks. The quantified fluorescent intensity or cell count at each discrete time point were respectively normalized by the averaged intensity or cell count from the ipsilateral side of the *zero distance* sections of the sham mice.

For neurogranin, in addition to the overall signal intensity, we also quantified its relative intensity in the soma to infer its cellular translocation (FI_soma_). Sections within 0.1 mm anterior-posterior to the optic fiber were selected for this analysis. It was easy to identify the neurogranin-positive neurons for sections from all three sham groups and the two stimulated groups, respectively, from 90 min-post and 48 h-post stimulation. Signals from these neuronal somas are often clearly organized in round shapes with brighter intensity against the salt-and-pepper pattern in the background. In contrast, sometimes the soma intensity is darker for the stimulated group at 24 h-post stimulation. To select the neurons without a bias, we firstly checked if any round cell-like shape is detectable by zooming into the micrographs. We further examined all round-circled signal clusters: soma area with signal dots were included, while empty somas were excluded to rule out non-pyramidal neurons. The selected somas were also verified with the channel-merged micrograph of YFP signals. Since YFP was sparsely tagged with pyramidal neurons, we finally noted down the soma size for post-hoc examination. In all the identified neurons, we first drew the soma shape and measured its fluorescent signal intensity and area size. Then we moved the mask to the neighboring area around the soma and measured the fluorescent intensity of the same-sized area as a reference. Five random selections were measured in the adjacent regions and averaged to serve as the reference. We normalized the signal intensity of soma by the signal intensity of its neighboring area as the relative soma intensity. We found that soma size and soma signal intensity were uncorrelated, so all the data points were used for statistical analysis.

### Statistical analysis

We have different types of data in the current study, non-clustered independent data and nested data. Independent measurement, such as behavioral data, was contributed only once by each mouse. We used the non-parametric Mann Whitney U test to examine if the optogenetic stimulation triggered significant behavioral alterations.

Some datasets, such as the signal intensity of immunohistochemical staining and the neural morphology, are highly nested. In the staining experiments, each mouse contributed multiple brain sections in five distance groups; for the morphological data, each mouse contributed several neurons, and each neuron further contributed multiple dendrite segments. In such conditions, using multiple measurements from each mouse as independent measurements artificially inflates sample size *N* and risks our study for achieving inappropriate conclusions. Indeed, we observed highly significant results for all quantified data obtained in IHC and microinjection experiments, when applied tests such as Mann Whitney U test or Kruskal Wallis test (Supplementary Materials: Comparing LMM with other tests). We therefore used a more conservative analysis, linear mixed effects model (LMM) in R (R Core Team 2019), to assess the effects of stimulation, while accounting for the nested residual structure. We used the lme4 package (Bates et al. 2015) and applied glmer function (GLMM) to model cell counting data. Our null hypothesis is that there is no significant difference between the sham and the stimulated mice, neither between the ipsi- and contralateral hemisphere of each mice. So in the model, we set the main effects of stimulation, implantation site, and their interaction effects as fixed effects. On the other hand, neuron ID, animal ID, and distance level are random effects (Zuur et al. 2009). Three discrete time points were separately analyzed. All models were checked in terms of homogeneous and normally distributed residuals, using diagnostic plots. We further checked final models for over-dispersion. Detailed model structures and the model selection and validation processes were described in the Supplementary Materials. All the R scripts of LMM and GLMM could be found under the following link: https://github.com/ErbB4/LMM-GLMM-R-plasticity-paper

The significance of fixed effects was tested by extracting effect strengths of each parameter, including their confidence intervals; *p <* 0.05, *p <* 0.01, and *p <* 0.001 were used to indicate 95% CI, 99% CI, or 99.9% CI of the estimated coefficient does not cross zero. If not stated otherwise, *∗* denoted the main effect of optogenetic stimulation (sham/stimulated), # denoted the main effect of stimulation side (ipsi/contra to the optic fiber). Significant interaction effects were not denoted but stated in the main text. “n.s.” denoted neither main effects nor interaction effects were significant.

### Neuron, synapse, and network models

Numerical simulations of networks with homeostatic structural plasticity were used as a framework to interpret the outcome of our various measurements in mouse experiments. We used the same neuron model, synapse model, and network architecture, as published in our previous paper on transcranial electric stimulation (Lu et al. 2019). All the plastic neuronal network simulations of the current study were performed with the NEST simulator using a MPI-based parallel configuration (Linssen et al. 2018).

The current-based leaky integrate-and-fire (LIF) neuron model was used for both excitatory and inhibitory neurons. We employed an inhibition-dominated recurrent network with 10 000 excitatory and 2 500 inhibitory neurons to represent the local network of ACC (Brunel 2000). All neurons in the network receive Poisson drive at a rate of *r*_ext_ = 30 kHz to reflect external inputs. All connections involving inhibitory neurons in this network were established randomly with 10% connection probability and then kept fixed. Only excitatory to excitatory (E-E) connections were subject to homeostatic structural plasticity (HSP) (Gallinaro and Rotter 2018; Lu et al. 2019). Each excitatory neuron monitors its own firing rate using its intracellular calcium concentration and grows or retracts its spines and boutons to form or break synapses (Figure 6A, see Supplementary Materials for details). Initially, the network has no E-E connections at all, but spontaneous growth goes on until an equilibrium between network activity and structure is reached. In this equilibrium state, the E-E connectivity is at 9%, and all excitatory neurons fire around the rate of 8 Hz imposed by the controller. Spiking activity is generally asynchronous and irregular. Detailed parameters of the neuron model, synapse model, and network model can be found in Supplementary Tables 4, Table 5, and Table 6. The methods employed to measure neuronal activity and connectivity in numerical experiments are described again in more detail in the Supplementary Materials.

**Figure 6:**
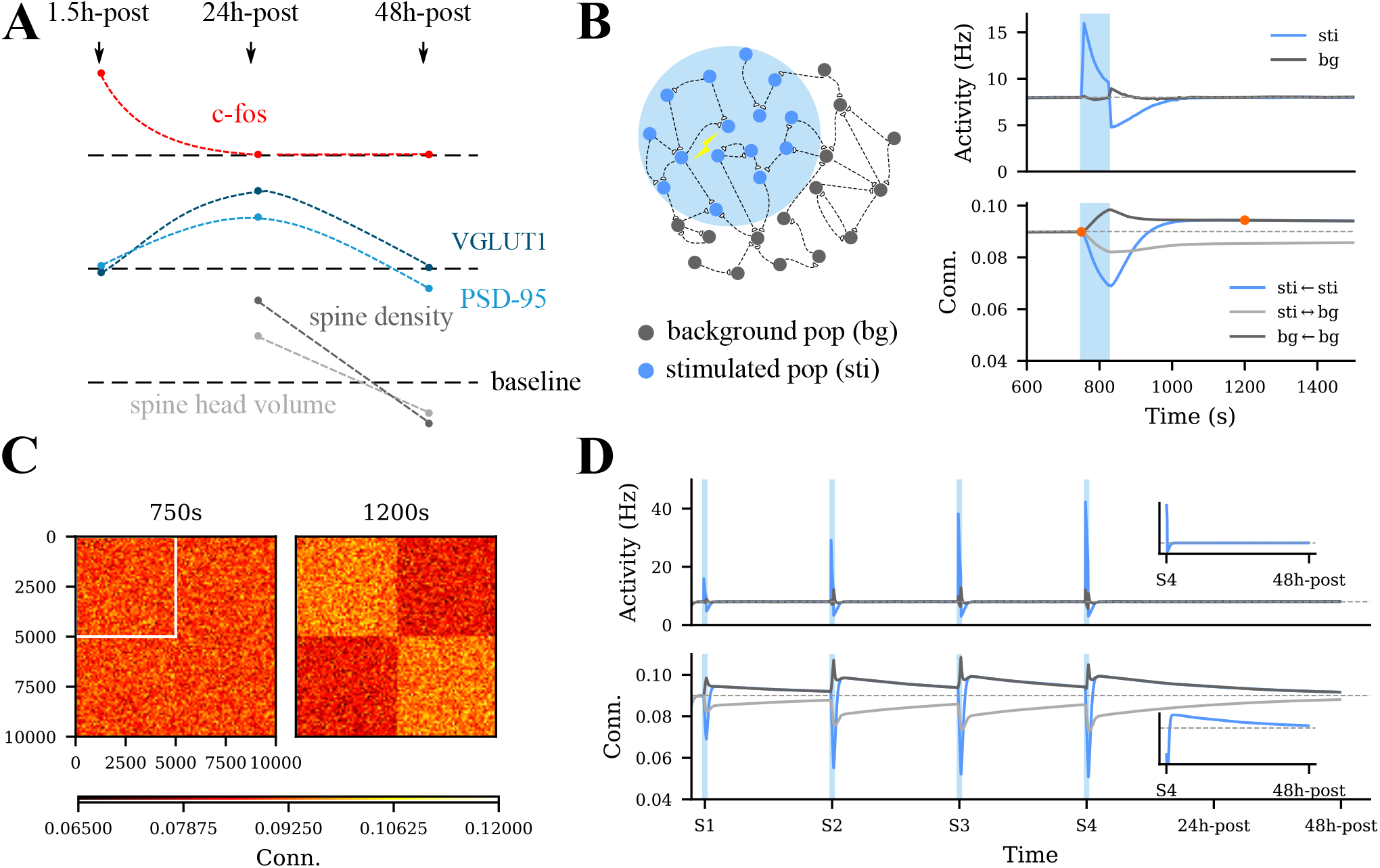
The homeostatic structural plasticity model. **A** The interpolated time course of the expression of c-Fos, VGLUT1, PSD-95 and spine morphology compared to sham (baseline). **B** Temporal evolution of neural activity and network connectivity in response to optogenetic stimulation. Light blue shaded areas indicate the stimulation period. Blue and dark gray curves in the upper panel represent the firing rate of stimulated and non-stimulated neurons. Blue, dark gray, and light gray curves in the lower panel represent the synaptic connectivity within or between populations. The blue and light gray curves finally coincided due to identical population sizes. **C** The connection matrix of all excitatory neurons at two time points before and after stimulation, labeled by orange dots in panel **B**. Columns are the presynaptic neurons, and rows are the postsynaptic neurons. Color indicates the average connectivity. The white square labels the intra-group connectivity of the stimulated neurons. **D** Repetitive optogenetic stimulation boosted the connectivity among the stimulated neurons. Small insets display the dynamics within 48 h after the last stimulation session.

### Modeling optogenetics

Optogenetics uses microbial opsin genes to achieve optical control of action potentials in specific neuron populations (Yizhar et al. 2011). In our current study, we used humanized channelrhodopsin-2 (hChR2), a fast light-gated cation-selective channel, to depolarize mouse pyramidal neurons (Nagel et al. 2003). The kinetics of ChR2 activation is a complicated light-dark adaptation process: Light activates and desensitizes the channels, while they recover in the dark phase (Bruun et al. 2015; Zamani et al. 2017). These state transitions have been studied in detail in computational models of ChR2 (Nikolic et al. 2013; Williams et al. 2013). It did not seem necessary, however, to include the detailed kinetics of ChR2 in our large spiking neural network with homeostatic structural plasticity. To reduce the complexity of the model and save computational power, we conceived optogenetic stimulation as an extra Poisson input of rate *r*_opto_ = 1.5 kHz and weight *J*_opto_ = 0.1 mV. Neurons responded with an increased firing rate to this stimulation, as observed in optogenetic stimulation experiments.

### Numerical experimental design

In mouse experiments, the ACC of animals were optogenetically stimulated for four consecutive days at the same time, with a duration of 30 min per day. In our computational model, we started the optogenetic stimulation after the network had reached its structural equilibrium. To avoid excessively long simulation time, we accelerated the remodeling process by employing relatively fast spine and bouton growth rates, see Figure 2 of Gallinaro and Rotter (2018) for details. The relative duration of stimulation vs. relaxation was left unchanged, however. Since the optogenetic stimulation in experiments activates a large fraction of all pyramidal neurons in ACC, we chose to stimulate half of the excitatory neurons (*f*_opto_ = 50 %) in the model. All the model parameters are summarized in Supplementary Table 7.

**Figure 2:**
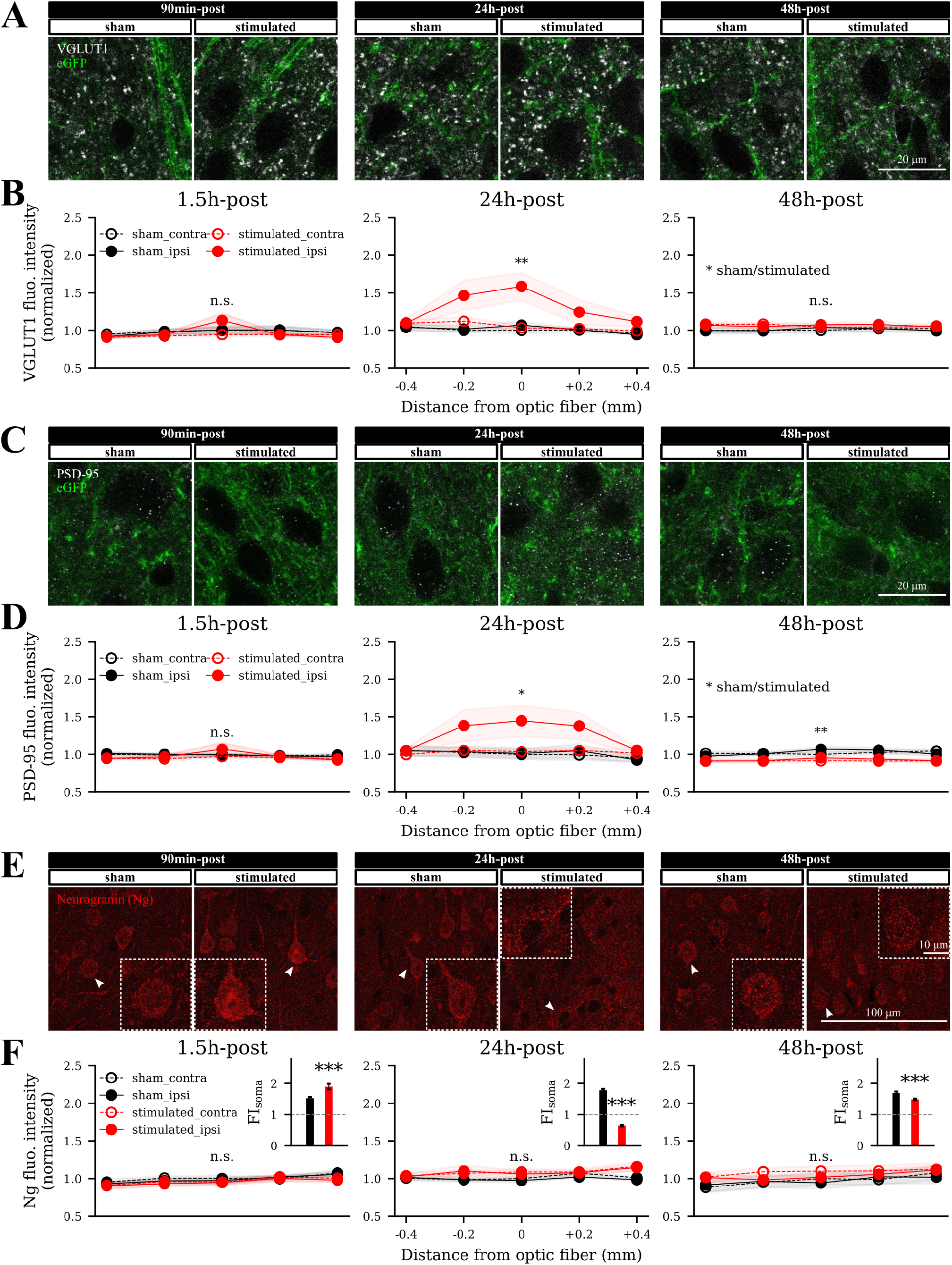
Optogenetic stimulation altered the expression of VGLUT1 and PSD-95 and the cellular translocation of neurogranin. **A, C, E** Representative images of VGLUT1, PSD-95, and neurogranin staining on the ipsilateral hemisphere of sections within 0.1 mm anterior-posterior (AP) to the optic fiber from both sham and stimulated mice. The insets in panel **E** display the example neurons. **B** Normalized VGLUT1 fluorescent intensity at different times and different distances to the fiber optic. The main effect of optogenetic stimulation (sham/stimulated) was significant at 24 h. **D** The PSD-95 fluorescent intensity at different times and distances. At 24 h and 48 h, the main effect of stimulation was significant. **F** The overall fluorescent intensity of neurogranin was not changed by stimulation (LMM), while the relative intensity of neural soma (FI_soma_) from the stimulated mice was slightly increased at 1.5 h, greatly dropped below 1 at 24 h, and recovered to a level lower than sham at 48 h (*p <* 0.001, *p <* 0.001, *p <* 0.001, respectively, Mann Whitney U test).

## Results

### Optogenetic activation of ACC (24a/24b) pyramidal neurons triggered cortical hyperactivity and behavioral alterations

To study the time course of neural structural plasticity, we adopted the optogenetic mouse model previously published in our laboratory (Barthas et al. 2015, 2017) in which we repetitively activated ACC pyramidal neurons for four days. Using electrophysiological recordings and optogenetic approaches, we have previously shown that the hyperactivity of anterior cingulate cortex drives chronic pain-induced anxiodepressive-like behavior. Therefore, we wondered whether activating ACC pyramidal neurons via optogenetic stimulation can trigger the same molecular and behavioral changes as seen in depressive-like behaviour. Indeed, the functional validation of ChR2-YFP expression using *ex vivo* electrophysiological recordings confirmed that the optogenetic stimulation at 20 Hz reliably triggers action potential firing of the ACC pyramidal neurons. We also observed *in vivo* that ACC optogenetic stimulation leads to a robust induction of c-Fos, compared to controls. In addition, four consecutive days of stimulation (30 minutes per day) were sufficient to trigger an MPK-1 upregulation in the ACC, as well as depressive-like phenotypes in naive animals (Barthas et al. 2017). Although a single stimulation might have been sufficient to trigger structural plasticity, in our structural plasticity analyses we focused on the effect of chronic optogenetic stimulation to induce robust and reproducible behavioural and molecular alterations.

First, we compared the viral transfection and transgenic approaches (Supplementary Figure 1-1A-E). We have previously shown with the transgenic approach that there was an increased c-Fos expression at 1.5 h after the stimulation (Barthas et al. 2015) and here we also reproduced the results with viral injection approach (Supplementary Figure 1-1F-G; Figure 1G, left panel). Besides the cortical hyperactivity, both approaches induced a depressive-like phenotype in mice at 24 h and 48 h post-stimulation (Supplementary Figure 1-1H-I; Figure 1E-F) as published before. To avoid double surgeries, we decided to continue with transgenic mice throughout the current study. We also confirmed that light did not trigger behavioral alterations (Supplementary Figure 1-2).

**Figure 1:**
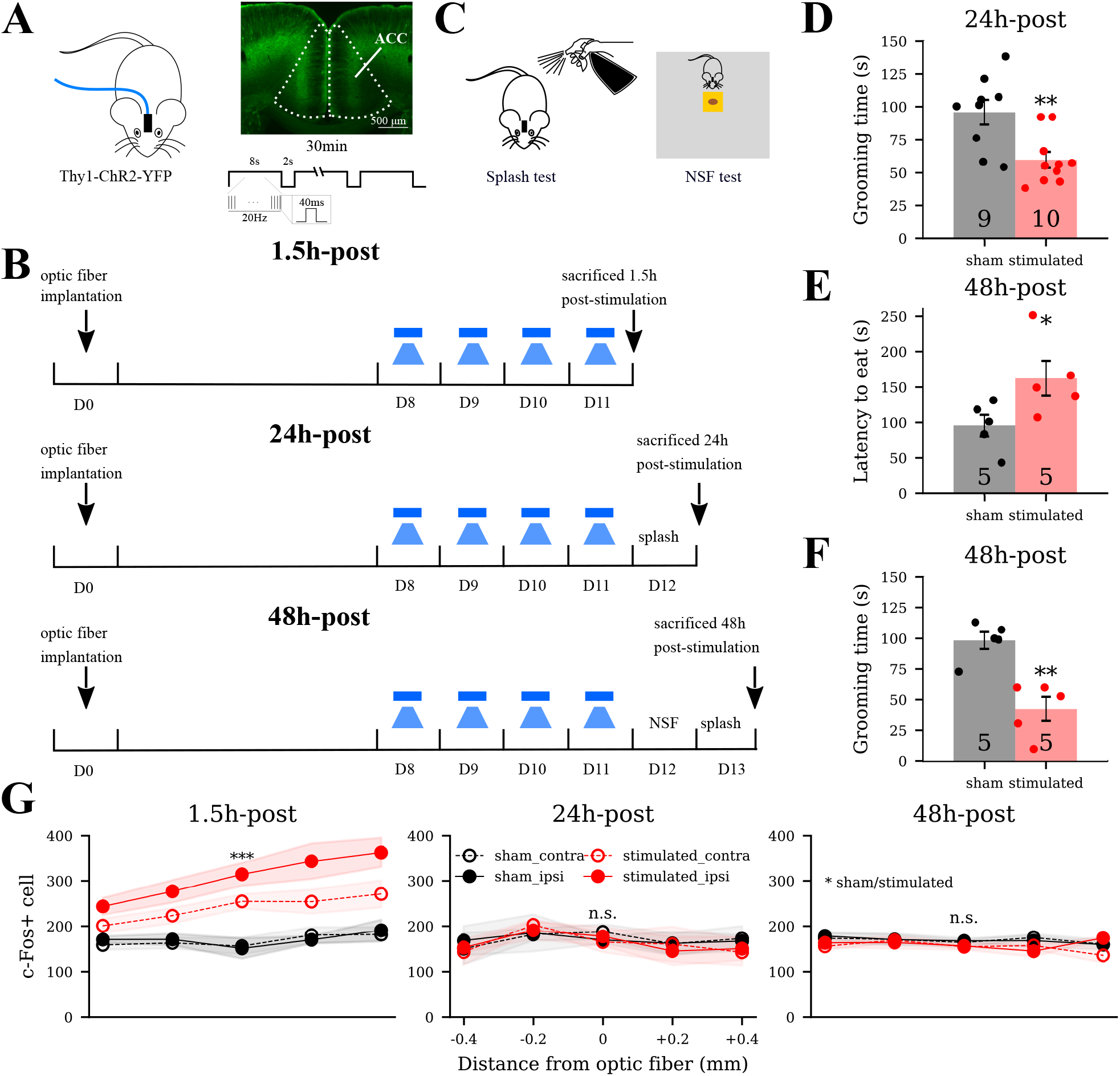
Consecutive optogenetic stimulation triggered hyperactivity in ACC and depressive-like behavior in mice. **A-C** Experimental design of the current study. **D-F** Mice used for immunohistochemical staining experiments showed depressive-like behavior at 24 h and 48 h post-stimulation. **D** Splash test results of mice sacrificed at 24 h post-stimulation (*p* = 0.003, Mann Whitney U test; *N* = 9 for sham, *N* = 10 for stimulated). **E-F** Results of NSF and splash tests for mice sacrificed at 48 h (*p* = 0.01 and *p* = 0.005, Mann Whitney U test; *N* = 5 for sham, *N* = 5 for stimulated mice). **G** c-Fos expression was elevated by the optogenetic stimulation at 1.5 h (*p <* 0.001) and specifically on the ipsilateral side (*p <* 0.01), decayed to baseline at 24 h and 48 h post-stimulation. The x ticks label the mean distance of stained sections (−, anterior; +, posterior) from the optical fiber. The y axis labels the c-Fos+ cell number within the masks of stained sections. GLMM was used for the statistical analysis. *p <* 0.05, *p <* 0.01, and *p <* 0.001 means 95% confidence interval (CI), 99% CI, and 99.9% CI does not cross zero respectively.

In two other batches of transgenic mice, we also examined the c-Fos expression at 24 h and 48 h post-stimulation and observed no difference between the stimulated and sham mice (Figure 1G, middle and right panels). These data suggested that optogenetic stimulation triggered hyperactivity in the ACC was restored to baseline level at 24 h and 48 h post-stimulation.

To capture the temporal evolution of neural plasticity, we stimulated transgenic Thy1-ChR2-YFP male adult mice and harvested their brains at 1.5 h, 24 h, and 48 h post-stimulation for further experiments (Figure 1A-C). Mice used for immunohistochemical staining experiments showed depressive-like behaviors at 24 h and 48 h post-stimulation, as shown by decreased grooming behaviour in splash test and increased latency to eat in the NSF test (Figure 1D-F). All mice group information and experimental design were summarized in Supplementary Table 1.

### VGLUT1 and PSD-95 in ACC showed time-dependent regulation by optogenetic stimulation

Structural plasticity could be reflected in changes of the number or size of synaptic contacts, but also in changes of network connectivity. Quantifying changes in network connectivity, however, is an unsolved challenge in the field. Before zooming directly into the microscopic analysis of synaptic contacts, using immunohistochemical staining against synaptic proteins, we determined when and where structural plasticity occured. We first quantified the overall expression of presynaptic VGLUT1 and postsynaptic PSD-95 in the ipsi- and contralateral hemispheres of ACC sections with reference to the hemisphere where the optic fiber was implanted, at 1.5 h, 24 h, and 48 h (Supplementary Table 1). Frontal-sectioned brain slices in both sham and the stimulated mice were organized based on their distance away from the optic fiber.

The representative fluorescent staining of VGLUT1 was organized by distance and by time in Supplementary Figure 2-1. Intensity quantification summarized in Figure 2A-B showed that optogenetic stimulation did not trigger significant alteration at 1.5 h (95% CI = [*−*0.083, 0.070], LMM), while significant upregulation was observed in the stimulated mice compared to sham mice at 24 h (99% CI = [0.009, 0.477], LMM). Further examination of interaction effects confirmed stronger upregulation in the ipsilateral side (99.5% CI = [*−*0.344*, −*0.116], LMM) in the stimulated mice. At 48 h, no more significant difference was detected between the stimulated and sham mice (95% CI = [*−*0.012, 0.138], LMM). Our data at discrete time points suggested that optogenetic stimulation altered VGLUT1 expression in a time-dependent manner. Indeed, the upregulation was observed after 1.5 h, peaked around 24 h, and returned to baseline at 48 h post-stimulation, while the stimulation effects were constrained to areas around the optic fiber.

Similar expression pattern was observed with PSD-95 staining (Supplementary Figure 2-2, Figure 2C-D). At 1.5 h, no significant changes were induced by the stimulation (95% CI = [*−*0.051, 0.018], LMM). At 24 h, enhanced expression of PSD-95 in ACC was observed in the stimulated mice (95% CI = [0.008, 0.55], LMM) and specifically in the ipsilateral hemisphere (99.5%CI = [*−*0.335*, −*0.109], LMM). At 48 h, although the effect size was small, the PSD-95 expression of the stimulated mice declined to a lower level than sham (99% CI = [*−*0.171*, −*0.005], LMM). Our data suggested a similar time-dependent manner of PSD-95 upregulation as VGLUT1 after the optogenetic stimulation: upregulation at 24 h and downregulation at 48 h.

### Neurogranin was not upregulated by optogenetic stimulation

Since PSD-95 is expressed in the postsynaptic membrane of glutamatergic synapses in both excitatory and inhibitory neurons (Zhang et al. 1999), we studied another post-synaptic protein, neurogranin, which is exclusively expressed in the pyramidal neurons (Singec et al. 2004). Despite the fact that the same type of quantification and analysis procedures were applied to neurogranin stained ACC sections, no time-dependent or side-dependent alterations of neurogranin was observed (Supplementary Figure 2-3, Figure 2E-F; 95% CI = [*−*0.111, 0.031], 95% CI = [*−*0.170, 0.277], 95% CI = [*−*0.102, 0.266], respectively, LMM).

Considering that neuronal stimulation could drive the translocation of neurogranin from soma to dendrites (Huang et al. 2011), we suspected that the optogenetic stimulation might fail to trigger neurogranin upregulation but induced the cellular translocation. Consequently, we selected sections within 0.1 mm anterior-posterior to the optic fiber and quantified the relative fluorescent intensity of neurogranin in the soma (insets in Figure 2E-F). Pyramidal neurons from the three sham groups all showed a high soma concentration. After the stimulation, the relative signal intensity of soma was slightly increased at 1.5 h (*p <* 0.001, Mann Whitney U test), decreased to a level lower than 1 at 24 h (*p <* 0.001, Mann Whitney U test), and recovered to a level above 1 but lower than the sham group at 48 h (*p <* 0.001, Mann Whitney U test). These data suggested that the optogenetic stimulation may not trigger neurogranin upregulation, but induce translocation with time: concentrated in soma at 1.5 h, translocated away from soma at 24 h, and recovered at 48 h. Since neurogranin is essential for the calcium concentration dependent translocation of calmodulin (Huang et al. 2011), the translocation of neurogranin hinted at the fact that calcium based plastic changes mostly occur in non-somatic compartments, most probably the dendrites.

### Dendritic tree structure was not drastically affected by optogenetic stimulation at **24 h** and **48 h**

The analysis of the expression of VGLUT1, PSD-95, and neurogranin suggest dendritic changes on the ipsilateral side in sections close to the optic fiber from the stimulated mice, at 24 h and 48 h after the stimulation. The amount of VGLUT1 and PSD-95 correlates with the total synaptic strength for all synapses. To specify whether the change of synaptic proteins corresponds to an alteration of synapse numbers or modifications of individual synaptic strength, we injected a fluorescent dye to visualize and analyze the neuronal morphology.

We then stimulated mice the same way as described above and harvested their brains at 24 h or 48 h post-stimulation. As shown in Figure 3A, mice showed depressive-like behavior as expected (*p* = 0.004 for 24 h-post group, *p* = 0.012 and *p* = 0.006 for 48 h-post group, Mann Whitney U test). We injected red fluorescent dye (Alexa 568) into pyramidal neurons selected from the area around the optic fiber to visualize and analyze the neuronal morphology.

**Figure 3:**
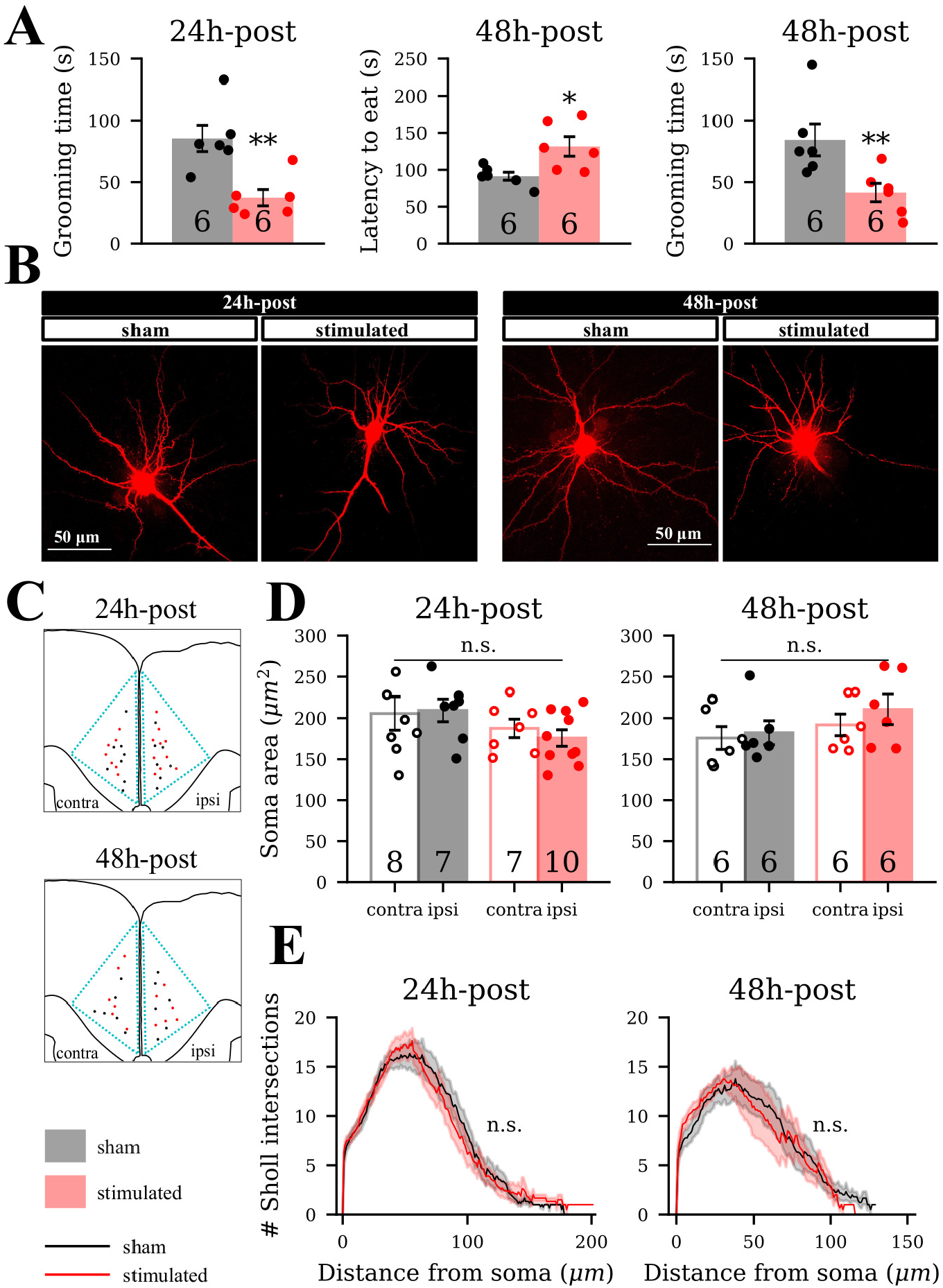
Neuronal dendritic tree structure was not drastically affected by the optogenetic stimulation at 24 h and 48 h. **A** Mice sacrificed at 24 h and 48 h for microinjection experiments showed depressive-like behavior in splash and NSF tests (for both batches, *N* = 6 for sham and *N* = 6 for the stimulated; *p* = 0.006, *p* = 0.012, and *p* = 0.004 respectively, Mann Whitney U test). **B** Representative example of neurons filled with red fluorescent dye. **C** Overall distribution of pyramidal neurons injected in layer 2-3 of ACC from both hemispheres for both batches. Black dots are from sham mice, and red dots are from the stimulated mice. For the 24 h-post group, we selected 32 well-injected neurons in total and 15 neurons were from sham mice; each mouse contributed 2.67 neurons on average (SD = 1.31). For the 48 h-post group, we selected 24 neurons in total and 12 neurons were from sham mice; each mouse contributed 2 neurons on average (SD = 2.12). **D** The soma size was not changed by stimulation (LMM). **E** Dendritic tree structure was not altered by stimulation (GLMM).

Neural dendritic structure at 24 h and 48 h was visualized as in Figure 3B. Pyramidal neurons from both ipsi- and contralateral ACC were collected (Figure 3C). The soma size and dendritic tree structure evaluated by Sholl intersections were not changed by the optogenetic stimulation (Figure 3D-E). No remarkable changes were detected in neither dendritic length nor average dendritic diameter, except that some dendritic segments showed a reduction or increase in dendritic diameter (Supplementary Figure 3-1). Our data suggested, except for local dendrite diameter changes, no drastic dendritic tree structure and soma size inflation or shrinkage of pyramidal neurons in the vicinity of the optic fiber were induced by the optogenetic stimulation.

### Optogenetic stimulation induced the opposite spine morphological changes at **24 h** and **48 h**

To further analyze morphological changes at dendritic level, we sampled several secondary to third level apical and basal dendritic segments from each neuron and did the 3D reconstruction of spines (Figure 4A-B). Besides spine density, we also evaluated the spine head volume and classified different types of spines such as filopodia, long-thin, stubby, and mushroom.

**Figure 4:**
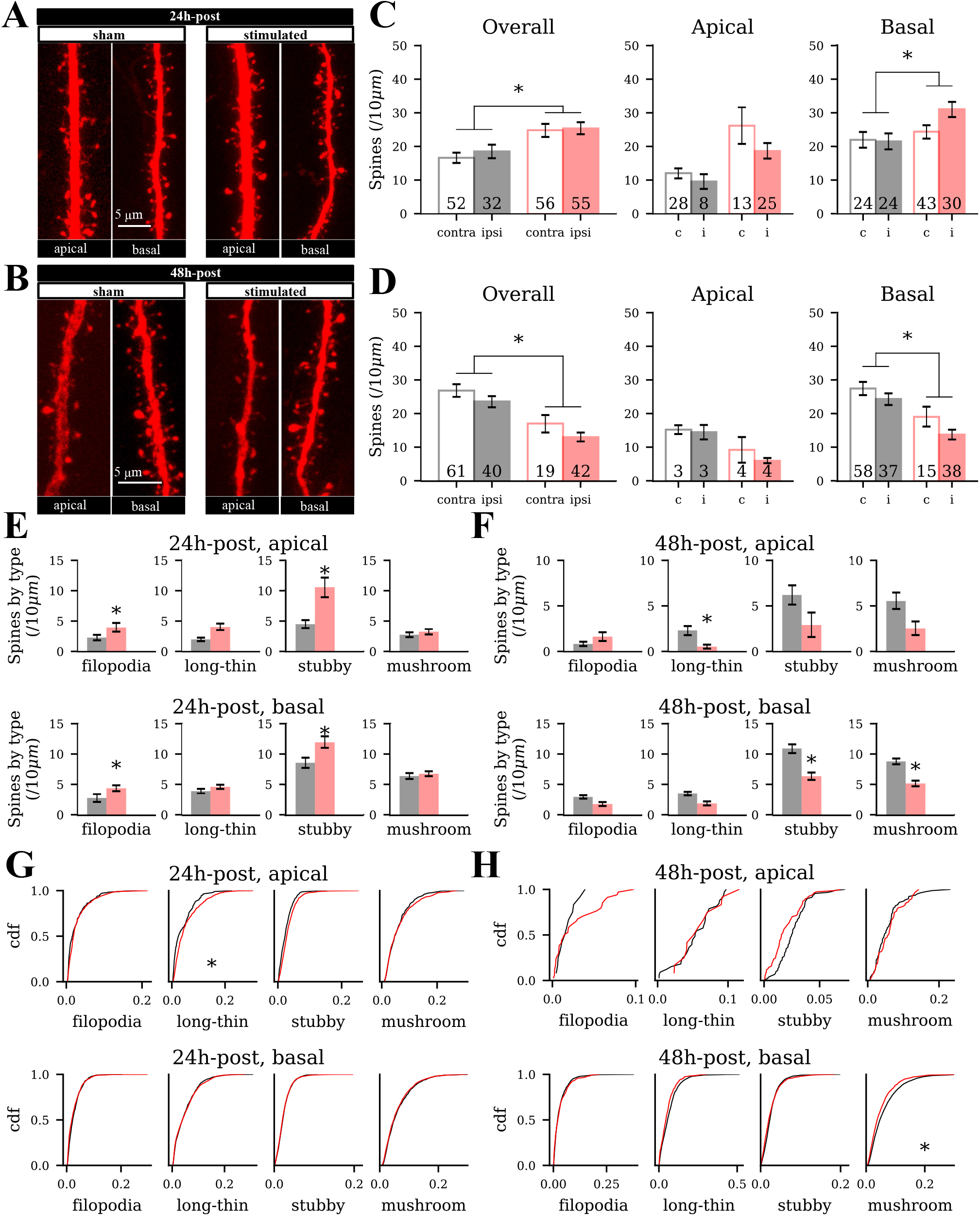
Spine density and head volume showed the opposite changes at 24 h and 48 h post-stimulation. **A-B** Representative example of filled dendritic segments. **C-D** Spine density at 24 h (84 dendritic segments from sham and 111 segments from stimulated mice) and 48 h (101 segments from sham and 61 segments from stimulated mice). **E-F** Spine density of each class. **G-H** Cumulative distribution of spine head volume. LMM was used for statistical analysis.

As shown in Figure 4C-D, the overall spine density was increased at 24 h (*p <* 0.05, LMM) and decreased at 48 h (*p <* 0.05, LMM) post-stimulation. The apical spine density showed the same tendency but the changes were not statistically significant; the basal dendrites showed significant spine density alterations (*p <* 0.05 and *p <* 0.05, LMM). Analysis by spine type suggested subtle changes in different spine types (Figure 4E-F). At 24 h, the spine density of filopodia and stubby type was increased in both apical (*p <* 0.05 and *p <* 0.05 respectively, LMM) and basal dendrites (*p <* 0.05 and *p <* 0.05, LMM). At 48 h, the spine density of long-thin type was reduced in apical dendrites (*p <* 0.05, LMM), while the stubby and mushroom type were reduced in basal dendrites (*p <* 0.05 and *p <* 0.05 respectively, LMM). These data suggested optogenetic stimulation triggered spinogenesis and spine retraction in both apical and basal dendrites at 24 h and 48 h post-stimulation respectively.

In addition, spine head volume data (Figure 4G-H) showed different regulation in apical and basal dendrites. At 24 h, the head volume distribution of long-thin spines in the apical dendrites was right-shifted to larger mean values by the optogenetic stimulation (*p <* 0.05, LMM) while no changes were detected in the head volume of basal dendrite spines. At 48 h, the spine head volume of mushroom spines in basal dendrites was left-shifted to smaller mean values by the optogenetic stimulation (*p <* 0.05, LMM), whereas the apical dendrites showed no significant difference. Our data suggested in addition to spine density changes, optogenetic stimulation induced spine enlargement and shrinkage at 24 h and 48 h post-stimulation respectively. The changes of overall spine density and spine head volume align with the evolution of PSD-95.

### Glial responses were involved in homeostatic plasticity induced by the optogenetic stimulation

Although glial cells were often linked with pathological brain states involving inflammation and injury, their involvement in regular synaptic plasticity has been frequently highlighted in recent studies (Dissing-Olesen et al. 2014; Haydon and Nedergaard 2015; Weinhard et al. 2018). The time course during the evolution of structural plasticity, however, has not been reported before. We thus stained glial fibrillary acidic protein (GFAP) as the markers for activated astrocytes (Hol and Pekny 2015) and ionized calcium-binding adaptor molecule 1 (IBA1) for both inactive and active microglia (Ohsawa et al. 2004) at 90 min, 24 h, and 48 h after the stimulation.

The fluorescent staining of GFAP was organized by distance and by time in Supplementary Figure 5-1. Our statistics analysis showed optic fiber implantation triggered astrocytes reactivation in both sham and stimulated mice, but the stimulation further enhanced the reactivation in the ipsilateral hemisphere throughout 48 h post-stimulation (Figure 5A-B). Similar results were observed with IBA1 staining (Supplementary Figure 5-2) as optogenetic stimulation induced significant enhancement of IBA1 expression at 24 h and 48 h post-stimulation (*p <* 0.001 and *p <* 0.001, GLMM) in the ipsilateral hemisphere and in the sections close to the optic fiber (Figure 5C-D). Since IBA1 labels microglia regardless of its activation state, the upregulation of IBA1 suggested microglia proliferation. Our data suggest that optogenetic stimulation triggered simultaneous astrocytes reactivation and microglia proliferation, and they displayed a time course that is different from the expression of synaptic proteins and neural morphological changes.

**Figure 5:**
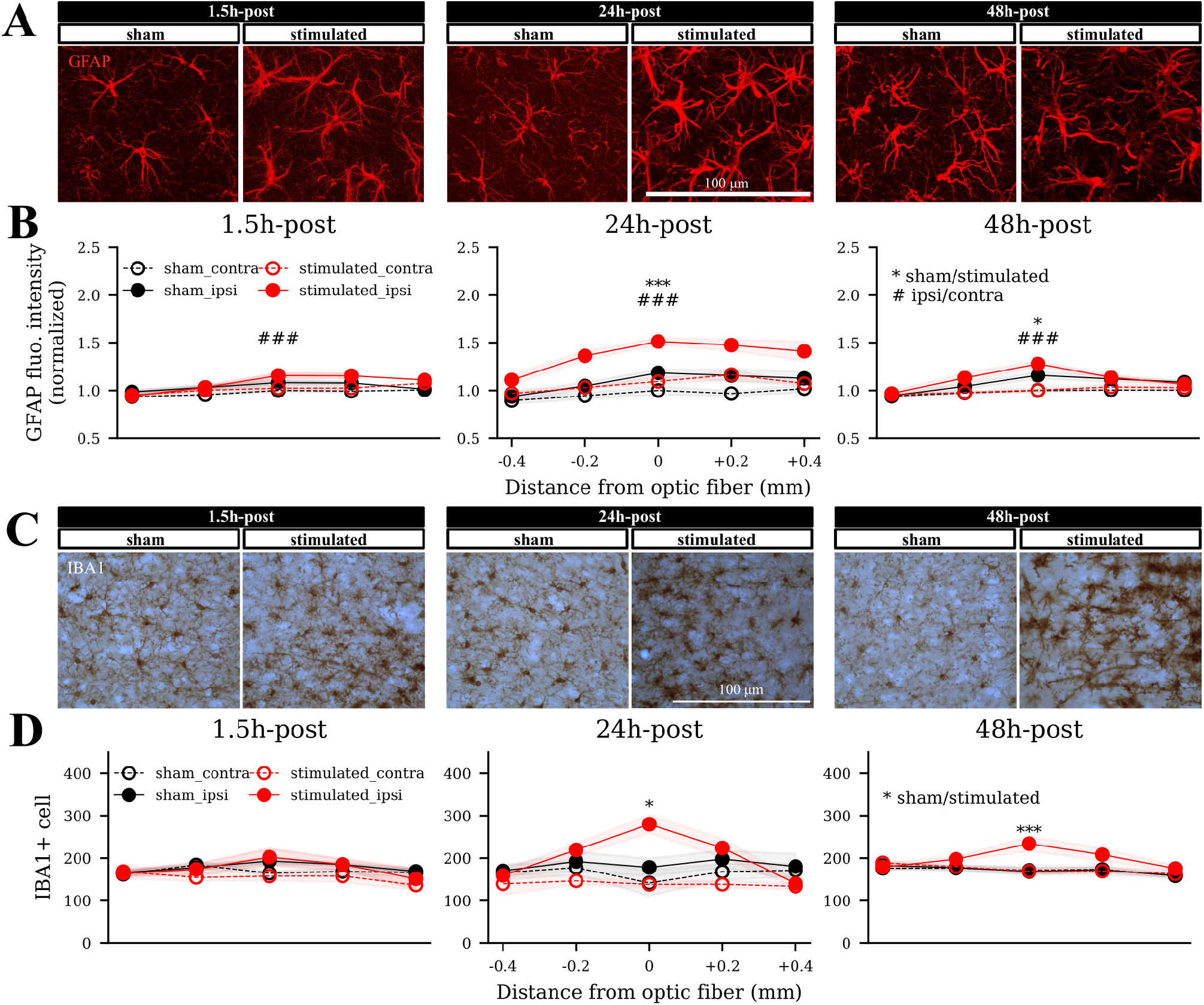
Expression of GFAP and IBA1 were upregulated by optogenetic stimulation at 24 h and 48 h post-stimulation. **A, C** Representative images of GFAP and IBA1 staining on the ipsilateral hemisphere of sections within 0.1 mm AP to the optic fiber from both sham and the stimulated mice. **B** The normalized GFAP fluorescent intensity at different time and distance. At 1.5 h, the main effects of side of stimulation were significant (99.5% CI = [0.0923, 0.0356], LMM). At 24 h, the main effects of optogenetic stimulation and side of stimulation were again significant (99.5% CI = [0.0646, 0.399], 99.5% CI = [0.182, 0.0628] respectively); their interaction effects were also significant (99.5% CI = [0.264, 0.109]). At 48 h, the main effects of stimulation, side of stimulation, and their interaction effects were significant (95% CI = [0.009, 0.094], 99.5% CI = [0.115, 0.043], 95% CI = [0.079, 0.006] respectively). **D** The IBA1+ cell counting at different time and distance to the optic fiber. At 1.5 h, only the interaction between stimulation and side of stimulation were significant (95% CI = [0.188, 0.00016], GLMM). At 24 h, the main effect of optogenetic stimulation was significant (95% CI = [0.025, 0.237]); their interaction effects were also significant (99.5% CI = [0.533, 0.105). At 48 h, the main effects of optogenetic stimulation was significant (99.5% CI = [0.073, 0.193]); their interaction was also significant (99.5% CI = [*−*0.208*, −*0.0369]).

### A computational model of stimulation-induced homeostatic structural plasticity

To achieve a clearer picture of the ongoing network remodeling dynamics in the current study, we interpolated the time course of synaptic protein expression and neural morphological changes within 48 h after four stimulation sessions (Figure 6A). The stimulation triggered immediate hyperactivity in ACC pyramidal neurons, as represented by c-Fos over-expression. Neural activity was restored to baseline at 24 h and 48 h. However, although neural activity was restored, we observed a delayed upregulation of synaptic proteins (VGLUT1 and PSD-95) and spine density at 24 h and a decrease to or below baseline at 48 h. These data suggest that the elevated synaptic proteins and spine density at 24 h do not contribute to sustaining high spontaneous neural activity. Three possibilities arise. (i) The upregulation of VGLUT1 and PSD-95 and spinogenesis failed to increase functional synaptic transmission among the stimulated pyramidal neurons. (ii) Optogenetic stimulation indeed increased the glutamatergic transmission, but additional mechanisms such as rapid E/I balance masked its effect on neural activity (Van Vreeswijk and Sompolinsky 1996; Shu et al. 2003; Zenke and Gerstner 2017). (iii) The upregulation of synaptic proteins and increase in spine number and volume are a consequence of firing rate homeostasis. Although we cannot directly reject options (i) and (ii) without electrophysiological recordings, it has indeed previously been shown that repetitive transcranial direct current stimulation (tDCS, 20 min *∗* 3 days) triggered enhanced synaptic transmission and increased spine density of pyramidal neurons 24 h after the stimulation in mice (Barbati et al. 2020). Besides, many previous studies have suggested that strength and morphology of excitatory synapses are homeostatically regulated (Turrigiano et al. 1998; Konur et al. 2003; De Gois et al. 2005; Ehrlich et al. 2007; Van Ooyen 2011) with or without changes of inhibitory synapses (Lenz et al. 2019; Knott et al. 2002) after activity perturbation. Therefore, it is highly possible that in our case optogenetic stimulation triggered homeostatic regulation.

The question now is which neuronal mechanism can account for the observed time course. Inhibitory STDP, inhibitory plasticity, synaptic scaling, and the Bienenstock-Cooper-Munro (BCM) model are commonly known homeostatic rules complementing Hebbian plasticity. These rules do not include synapse rewiring. In some cases, enhanced spontaneous neural activity emerges with enhanced synaptic weight, which does not fit what we observed at 24 h post-stimulation (Litwin-Kumar and Doiron 2014; Toyoizumi et al. 2014; Lazar et al. 2009; Zenke and Gerstner 2017). We thus selected the model of homeostatic structural plasticity (HSP), which assumes structural changes regulated by firing rate homeostasis. We simulated an inhibition-dominated spiking neural network to represent ACC (Figure 6B), in which optogenetic stimulation was introduced to half of the excitatory population. Transient stimulation perturbed the neural activity and triggered synapse turnover as a result of homeostatic structural plasticity (blue curves in Figure 6B), as described in a previous publication (Lu et al. 2019). As a result of synaptic reorganization, the connectivity among the stimulated neurons remained elevated after stable firing rates were achieved (Figure 6C). In a repetition protocol based on our *in vivo* experiments, simulation results show that the connectivity among the stimulated neurons increased after each repetition. Within 48 h after the final stimulation (S4), the firing rate of the stimulated neurons rapidly returned to baseline. The connectivity, however, remained elevated and decayed only slowly (insets in Figure 6D). Although we could not differentiate stimulated and non-stimulated neurons in the mouse experiments, the neurons selected for morphological analysis were close to the optic fiber and, therefore, had a higher chance to be stimulated. The HSP model, thus, provided a fully consistent explanation for the observed c-Fos expression and spine morphology.

## Discussion

In the current study, we combined both mouse experiments and computer simulations to study structural plasticity. We first systematically investigated the neural activation and neural morphology of the neocortical region anterior cingulate cortex (ACC) after chronic optogenetic stimulation in an *in vivo* mouse model. We found that the activation of a subset of excitatory neurons in ACC over four consecutive days triggered substantial alterations. In fact, the temporal profiles of specific molecular and morphological changes over 48 h post-stimulation were intertwined in a specific way. The expression of VGLUT1 and PSD-95, as well as the spine density and spine head volume, were above baseline at 24 h and restored to baseline or slightly below at 48 h. Intriguingly, although such changes seem to suggest altered synaptic transmission, neural activity estimated by c-Fos expression did not show any change at 24 h and 48 h. After neural activity has rapidly returned to baseline, synaptic protein expression and spine density undergoes a rise and a decay as compared to the control (Figure 7, red and blue curves). Altogether, it appears as if synaptic plasticity regulated by firing rate homeostasis can explain the time course of events described above quite well. In fact, we verified with the help of computer simulations that the homeostatic structural plasticity (HSP) model, in principle, recapitulates the observed biphasic dynamics (Gallinaro and Rotter 2018; Lu et al. 2019; Gallinaro et al. 2020).

**Figure 7:**
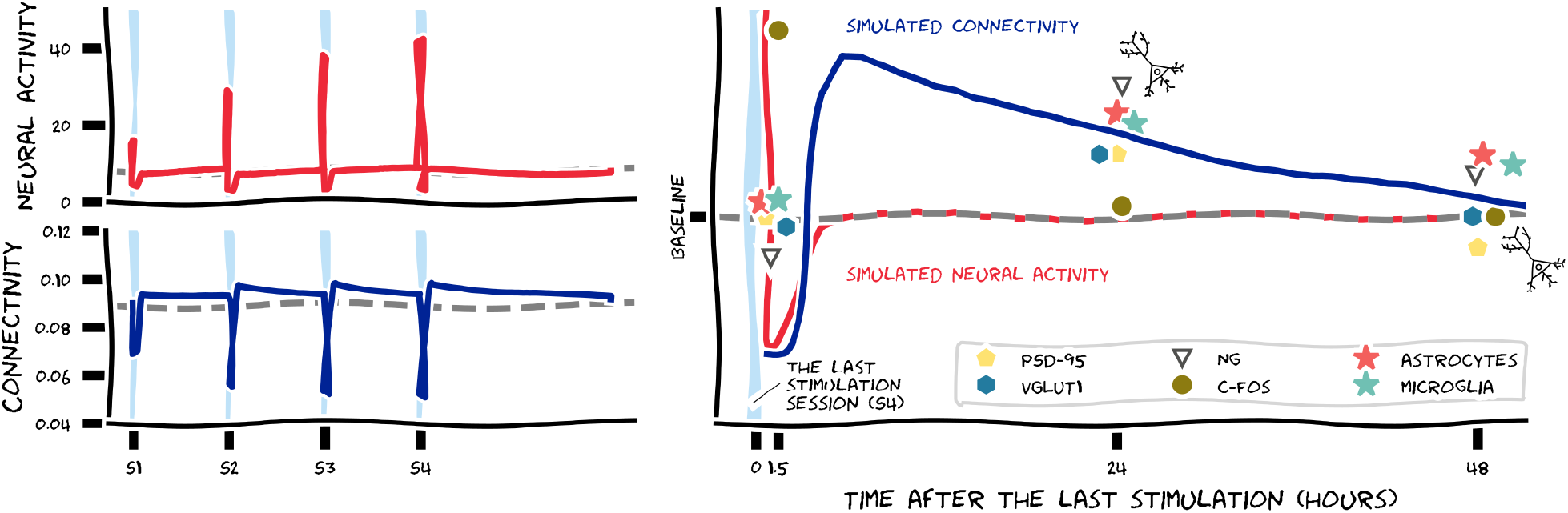
Joint summary of the results of mouse experiments and the computational model. Light blue shaded areas indicate four sessions of optogenetic stimulation (S1 to S4, 30 min per session). Red and blue curves represent the temporal profile of neural activity and connectivity of stimulated neurons, as predicted by the model. Colored symbols in the right panel represent the expression level of c-Fos, synaptic proteins, and glial markers, respectively. Measurements were made at 1.5 h, 24 h, and 48 h post-stimulation, shown are the values from stimulated mice relative to the corresponding sham group. For Neurogranin, we show its relative expression in dendrites. The gray dashed line represents the baseline. Symbols above or below the baseline indicate that values in stimulated mice are increased or decreased as compared to sham, respectively. The illustration of a neuron with dendrites and spines depicts that spine density and spine head volume were both increased at 24 h, but decreased at 48 h post-stimulation.

Our experiments elucidated how structural plasticity of pyramidal neurons evolves in time after optogenetic stimulation in the mouse experiments. Analysis of the expression of synaptic proteins clearly indicates that robust synaptic changes occurred at 24 h and 48 h after stimulation. VGLUT1 is the glutamate transporter protein that controls the quantal glutamate content of individual synaptic vesicles (Fremeau et al. 2004; Wilson et al. 2005). Therefore, the upregulation and downregulation of VGLUT1 observed in our current study hints at a change of glutamate release in synapses, corresponding to the presynaptic strength accumulated over many neurons in a given tissue volume. PSD-95 is a scaffold protein in the postsynaptic density which transforms rapidly (Gray et al. 2006) and which is actively redistributed between the synaptic sites and the cytoplasmic pool (Bresler et al. 2001; Gray et al. 2006). Functionally, PSD-95 organizes the distribution of AMPA receptors (Zhang et al. 1999; Chen et al. 2011) and modulates spine morphology (El-Husseini et al. 2000; Pak and Sheng 2003; Fossati et al. 2015). We observed similar biphasic changes in the expression of PSD-95 and VGLUT1 after stimulation.

However, the expression of neurogranin, a calmodulin-binding protein exclusively expressed in the soma and dendrites of excitatory neurons (Singec et al. 2004), behaved differently. Neurogranin has a molecular weight less than 10 kDa and thus can swiftly translocate within pyramidal neurons upon synaptic stimulation from the cell plasma to the nucleus (Garrido-Garćıa et al. 2009) or from the soma to the dendrites (Huang et al. 2011). We found that optogenetic stimulation failed to trigger upregulation or downregulation of neurogranin, but changed somatic signal intensity over time, which may suggest a translocation away from the soma in parallel with the upregulation of VGLUT1 and PSD-95. Functionally, neurogranin bidirectionally modulates synaptic plasticity by affecting the phosphorylation pattern of postsynaptic density proteins via calmodulin (Xia and Storm 2005; Zhong and Gerges 2012; Svirsky et al. 2020; Hwang et al. 2021). Upon activity perturbation, neurogranin also coordinates the conversion of AMPAR-silent synapses to AMPAR-active synapses, as well as synapse elimination (Han et al. 2017). Based on this evidence, we propose that translocated neurogranin may interact with postsynaptic density proteins. This could work in synergy with PSD-95 to modulate functional and plasticity. In addition, due to the dependence of translocation on calcium concentration (Huang et al. 2011) and the temporal profile of intracellular calcium dynamics, neurogranin might not be stored in spines and exert its impact on synaptic plasticity only at a later time.

All these observations point to a biphasic regulation of synaptic strength within 48 h and an orchestrated regulation of presynaptic and postsynaptic plasticity after stimulation, as reported before by others (Ehrlich et al. 2007; Letellier et al. 2019; Sanderson et al. 2020). The results of neural morphology analysis were in line with our observations in synaptic proteins. Optogenetic stimulation did not alter dendritic branching structure or soma size, as it was observed in the case of diseases (Chidambaram et al. 2019). Rather, stimulation induced biphasic changes at the level of dendritic spines. The density and volume of spines in the stimulated pyramidal neurons increased at 24 h and slightly decreased at 48 h, as compared to controls. This is highly interesting, as spine volume correlates with synapse strength (Matsuzaki et al. 2004) and PSD-95 clustering (Cane et al. 2014). Increased spine density increases the chances to form new synapses. All things considered, we conclude that synaptic transmission and connectivity is increased at 24 h and restored to baseline at 48 h after stimulation.

Given the biphasic temporal profile of changes in synaptic proteins and dendritic spine morphology, we hypothesized that they might be a result of homeostatic regulation. In fact, the time evolution of synaptic protein expression reflects the accumulation of effects of all four stimulation sessions. The time course of PSD-95, for example, is around baseline at 1.5 h, upregulated at 24 h, and decayed slightly below baseline at 48 h after the last stimulation. Classical Hebbian plasticity depends on positive feedback, and it would systematically increase PSD-95 expression upon stimulation. Homeostatic plasticity, in contrast, depends on negative feedback, and it would at least transiently decrease the PSD-95 level. As the low level of PSD-95 immediately after the fourth stimulation is also 24 h after the third stimulation, a pure Hebbian mechanism is out of question, and a contribution of homeostatic control is likely (see Supplementary Figure 6-1 for a graphical illustration of the argument). A second independent argument can be derived from the temporal profile of c-Fos expression. Indeed, the hyperactivity expected by optogenetic stimulation in pyramidal neurons is not visible 24 h and 48 h after stimulation, possibly because homeostatic regulation has brought it back to its set point. Although c-Fos expression is not a very accurate indicator of neural activity, other *in vivo* electrophysiological recordings did confirm the robust firing rate homeostasis in the cortex (Pacheco et al. 2019). As a result, it seems possible that the observed changes in synaptic proteins and spine morphology reflect the dynamic process of homeostatic control to restore neural activity after activity perturbation. Theoretically, an alternative explanatory scheme may be linked to inhibitory plasticity (Vogels et al. 2011, 2013), although we found no evidence for this in our data.

We proposed the model of homeostatic structural plasticity to explain the observed biphasic changes of spine morphology at 24 h and 48 h post-stimulation, when the neural activity indicated by c-Fos expression was already back at baseline. This model explains how neuronal firing rates are stabilized using structural plasticity linked with a homeostatic controller (Butz et al. 2009; Van Ooyen 2011; Butz and van Ooyen 2013). We have previously shown in computer simulations how external stimulation can trigger cell assembly formation by deleting connections and forming new synapses controlled by firing rate homeostasis (Gallinaro and Rotter 2018; Gallinaro et al. 2020; Lu et al. 2019). In computer simulations of the optogenetic experiment, we observed a very similar cell assembly formation process. Specifically, we showed that in this model the connectivity among stimulated neurons remained at a high level although the firing rate had already returned to baseline. Although we could not record the connectivity among the stimulated neurons in mouse ACC like we did in computer simulations, the changes of spine density in the pyramidal neurons sampled in the area close to the optic fiber served as a proxy and seemed to fit the computer simulations. Due to firing rate homeostasis, application and termination of the external stimulation should trigger slow homeostatic responses of opposite sign. Typical experiments, however, record a mix of changes occurring during perturbation and after perturbation, which may be of opposite sign. For instance, plastic changes observed as a result of a persistent lesion or denervation occur during input deprivation. In contrast, plastic alterations observed after stimulation, as in the current study, are mixed on and off effects. So it is critical to use an experimental design that includes both phases and measure during time periods that are long enough to re-establish the neural activity homeostasis. In addition to spine turnover and spine density, specific changes in connectivity represent another crucial feature that influences network function.

Nevertheless, the question arises whether similar effects on neural morphology would occur after a single stimulation already, rather than only after four stimulation sessions. Some alterations, such as developing a depressive-like phenotype, indeed requires chronic stimulation, as reported in our previous work (Barthas et al. 2017). The same rule, however, does not necessarily apply to changes in spine size and density. Chronic *in vivo* imaging has revealed ongoing spine turnover and fluctuation in spike head volume even under normal conditions (Holtmaat et al. 2006). However, external perturbation of neural activity can accelerate network remodeling considerably. The magnitude of such changes depends on circumstantial factors, such as the developmental stage, but also on stimulation parameters, including the type, intensity, and duration of the perturbation. Changes in spine head volume generally seem to come first. Both weak stimulation (e.g. using glutamate uncaging) and supra-threshold stimulation (e.g. using repetitive magnetic stimulation) were reported to induce considerable changes in spine volume within a few hours post-stimulation (Vlachos et al. 2012a; Matsuzaki et al. 2004; Noguchi et al. 2019). Longer and/or stronger stimulation is required to induce changes in spine density, which involves protein synthesis and *de novo* spinogenesis (Trachtenberg et al. 2002; Holtmaat et al. 2006). In experiments, the question arises whether a single stimulation can trigger neural morphological changes at all, and whether potentially subtle changes can be detected statistically. It is very well possible that changes in spine head volume occur already after a single stimulation, but they might be masked by the inevitable inter-neuron and inter-animal variance. Repetitive imaging of one and the same dendritic segment might be necessary to answer this question. Assuming that a single stimulation already triggers the structural remodeling process, we have shown in computer simulations in a previous study (Lu et al. 2019) and this manuscript that repetitive stimulation can be accumulated, increasing synaptic connectivity in an incremental way. This may also explain why, in our experiments, we could observe elevated c-Fos expression in the contralateral hemisphere of the stimulated mice, but no significant changes in neural morphology. Neural activation of the contralateral hemisphere could either be due to inter-hemispheric projections, or due to light penetrating from the ipsilateral side. Both effects are certainly weaker than the effects of direct stimulation. As already mentioned above, subtle changes triggered by weak stimulation might not be detectable by the analysis methods used in our current study.

Our study also casts light on the relation between ACC hyperactivity, synaptic plasticity, and depressive-like behavior. ACC is a hub for negative affects, pain, and their comorbidity (Humo et al. 2019), accompanied by different forms of synaptic plasticity (Bliss et al. 2016). Chronic pain can induce hyperactivity and synaptic potentiation in ACC, along with anxiodepressive behavior in mice (Sellmeijer et al. 2018; Koga et al. 2015). ACC hyperactivity artificially induced by optogenetic stimulation also generates depressive-like behavior in naive mice (Barthas et al. 2015, 2017). It is unclear, however, whether changes in neuronal activity, spine morphology, and depressive-like behavior develop in parallel due to a common condition, or whether there are causal links between individual factors (Gipson and Olive 2017). In our experiments, neural activity quickly decayed to baseline after stimulation, but the mice exhibited sustained depressive-like behavior, which can last for around two weeks after the stimulation was terminated (Barthas et al. 2017). So the alterations in depressive-like behavior seem to always lag behind changes in ACC neural activity. This evidence suggests that depressive-like behavior may be mediated by persistent changes which depend on the accumulated effects of neural activity. On the other hand, we observed that synaptic plasticity (synaptic proteins and spine morphology) exhibit transient changes during depressive behavior. This suggests that the time scales of structural plasticity and behaviour differ from each other.

Although glia cells are not the main focus of this study, we nevertheless showed that the time course of astrocyte activation and microglia proliferation matches the time course of homeostatic structural plasticity. Indeed, we observed increased numbers of IBA1- positive cells and enhanced GFAP expression throughout 48 h after stimulation, which suggests that both microglia cells and astrocytes were active at least between 1.5 h and 48 h post-stimulation. Microglia and astrocyte activation is often regarded as a sign of neural inflammation. This also raises the question whether the phenomena observed in our present study reflect a pathological state, or whether they should be interpreted as reflecting regular structural plasticity. We cannot fully exclude the possibility of neuroinflammation, as glial activation is heterogeneous, and there is no clear distinction between normal and pathological conditions anyway. Besides, accumulating evidence emphasizes the critical role of glial cells in cleaning up extracellular chemicals, fostering spinogenesis, and facilitating synaptic plasticity in normal brain development and plasticity (Eroglu and Barres 2010; Zamanian et al. 2012; Liddelow and Barres 2017). Glia-secreted proinflammatory cytokine tumor necrosis factor *α* (TNF *− α*), for instance, has been shown to also regulate synaptic transmission and homeostatic synaptic scaling (Stellwagen and Malenka 2006; Steinmetz and Turrigiano 2010). According to our data, the structure of dendritic trees and soma size were only weakly affected by optogenetic stimulation. Therefore, it seems unlikely that glial activation is caused by excitotoxicity and neural inflammation. Further studies are of course needed to explore if microglia and astrocyte activation triggered by optogenetic stimulation are involved in maintaining chemical homeostasis (Jo et al. 2014) and inducing morphological changes of spines (Weinhard et al. 2018).

In conclusion, our joint experimental-theoretical efforts provide evidence that, in response to supra-threshold optogenetic excitation, neurons modulate their synaptic connectivity to restore neural activity in a homeostatic way. The homeostatic structural plasticity model was able to qualitatively explain the observed time course of neural activity and spine morphology. Further joint work is needed to capture the effects of activity perturbation on specific network connectivity and include functional aspects to structural plasticity models.

## Supportive Information

### Funding information

This work was funded by the Universitätsklinikum Freiburg, University of Strasbourg, Centre National de la Recherche Scientifique (Grant/Award Number: UPR3212), NARSAD Young Investigator Grant from the Brain & Behavior Research Foundation (Grant/Award Number: 24736) and Young Talent Award from the University of Strasbourg (IdEx award). Additional funding was obtained by BrainLinks-BrainTools (funded by the Federal Ministry of Economics, Science and Arts of Baden-Württemberg within the sustainability program for projects of the excellence initiative II) and by the Carl Zeiss Stiftung. We are grateful for support by the state of Baden-Württemberg through bwHPC and the German Research Foundation (DFG) through grant no INST 39/963-1 FUGG (bwForCluster NEMO). HL is funded by a NEUREX fellowship and the Deutsch-Französische Hochschule.

## Author contributions

HL, JG, CN, SR, and IY conceived the project and designed the experiments. HL, JG, and SR established the computer model. HL performed the network simulation and analysis. HL performed all the mice experiments, collected data, and performed the analysis. CN and IY helped with the experiments. IY proposed the quantification strategy. HL wrote the manuscript, and all the authors revised and approved the paper.

## Acknowledgments

We thank Elisabeth Waltisperger and Alessandro Bilella for advice on immunohistochemical staining. We thank Stephan Doridot and the Chronobiotron animal facilities for breeding and genotyping the mice. We thank Beyza Ayazgok for the attempts in molecular analysis. We thank Josef Bischofberger, Stefan Vestring, and Cyril Bories for their help and advice in setting up the microinjection experiments. We also thank the staff of the Life Imaging Center (LIC) in the Center for Biological Systems Analysis (ZBSA) of Freiburg University, in particular A. Naumann and R. Nitschke, for help with their confocal microscopy resources and their excellent support in image recording and analysis. Advice and code review by Martin Mörsdorf concerning the statistical analysis using linear mixed models are sincerely acknowledged. The authors also thank Sylvain Hugel, Rudi Tong, and Sandra Diaz-Pier for useful discussions. We also thank Uwe Grauer from the Bernstein Center Freiburg, as well as Bernd Wiebelt and Michael Janczyk from the Freiburg University Computing Center for their assistance with HPC applications and data storage. The author HL is currently working in the Institute of Anatomy and Cell Biology at the University of Freiburg; the help and support from the Vlachos lab is sincerely acknowledged.

## References

Barbati SA, Cocco S, Longo V, Spinelli M, Gironi K, Mattera A, Paciello F, Colussi C, Podda MV, Grassi C. 2020. Enhancing plasticity mechanisms in the mouse motor cortex by anodal transcranial direct-current stimulation: the contribution of nitric oxide signaling. Cerebral Cortex. 30:2972–2985.

Barthas F, Humo M, Gilsbach R, Waltisperger E, Karatas M, Leman S, Hein L, Belzung C, Boutillier AL, Barrot M, et al. 2017. Cingulate overexpression of mitogen-activated protein kinase phosphatase-1 as a key factor for depression. Biological Psychiatry. 82:370–379.

Barthas F, Sellmeijer J, Hugel S, Waltisperger E, Barrot M, Yalcin I. 2015. The anterior cingulate cortex is a critical hub for pain-induced depression. Biological Psychiatry. 77:236–245.

Bates D, Mächler M, Bolker B, Walker S. 2015. Fitting linear mixed-effects models using lme4. Journal of Statistical Software. 67:1–48.

Bear MF, Malenka RC. 1994. Synaptic plasticity: LTP and LTD. Current Opinion in Neurobiology. 4:389–399.

Bi Gq, Poo Mm. 1998. Synaptic modifications in cultured hippocampal neurons: dependence on spike timing, synaptic strength, and postsynaptic cell type. Journal of Neuroscience. 18:10464–10472.

Bliss TV, Collingridge GL, Kaang BK, Zhuo M. 2016. Synaptic plasticity in the anterior cingulate cortex in acute and chronic pain. Nature Reviews Neuroscience. 17:485.

Bresler T, Ramati Y, Zamorano PL, Zhai R, Garner CC, Ziv NE. 2001. The dynamics of SAP90/PSD-95 recruitment to new synaptic junctions. Molecular and Cellular Neuroscience. 18:149–167.

Brunel N. 2000. Dynamics of sparsely connected networks of excitatory and inhibitory spiking neurons. Journal of Computational Neuroscience. 8:183–208.

Bruun S, Stoeppler D, Keidel A, Kuhlmann U, Luck M, Diehl A, Geiger MA, Woodmansee D, Trauner D, Hegemann P, et al. 2015. Light–dark adaptation of channelrhodopsin involves photoconversion between the all-trans and 13-cis retinal isomers. Biochemistry. 54:5389–5400.

Butz M, van Ooyen A. 2013. A simple rule for dendritic spine and axonal bouton formation can account for cortical reorganization after focal retinal lesions. PLOS Computational Biology. 9.

Butz M, Woergoetter F, van Ooyen A. 2009. Activity-dependent structural plasticity. Brain Research Reviews. 60:287–305.

Cane M, Maco B, Knott G, Holtmaat A. 2014. The relationship between PSD-95 clustering and spine stability in vivo. Journal of Neuroscience. 34:2075–2086.

Caroni P, Donato F, Muller D. 2012. Structural plasticity upon learning: regulation and functions. Nature Reviews Neuroscience. 13:478–490.

Chater TE, Goda Y. 2020. My neighbour hetero—deconstructing the mechanisms underlying heterosynaptic plasticity. Current Opinion in Neurobiology. 67:106–114.

Chen X, Nelson CD, Li X, Winters CA, Azzam R, Sousa AA, Leapman RD, Gainer H, Sheng M, Reese TS. 2011. PSD-95 is required to sustain the molecular organization of the postsynaptic density. Journal of Neuroscience. 31:6329–6338.

Chidambaram SB, Rathipriya A, Bolla SR, Bhat A, Ray B, Mahalakshmi AM, Manivasagam T, Thenmozhi AJ, Essa MM, Guillemin GJ, et al. 2019. Dendritic spines: revisiting the physiological role. Progress in Neuro-Psychopharmacology and Biological Psychiatry. 92:161–193.

De Gois S, Schäfer MKH, Defamie N, Chen C, Ricci A, Weihe E, Varoqui H, Erickson JD. 2005. Homeostatic scaling of vesicular glutamate and GABA transporter expression in rat neocortical circuits. Journal of Neuroscience. 25:7121–7133.

Diaz-Pier S, Naveau M, Butz-Ostendorf M, Morrison A. 2016. Automatic generation of connectivity for large-scale neuronal network models through structural plasticity. Frontiers in Neuroanatomy. 10:57.

Dissing-Olesen L, LeDue JM, Rungta RL, Hefendehl JK, Choi HB, MacVicar BA. 2014. Activation of neuronal NMDA receptors triggers transient ATP-mediated microglial process outgrowth. Journal of Neuroscience. 34:10511–10527.

Dumitriu D, Rodriguez A, Morrison JH. 2011. High-throughput, detailed, cell-specific neuroanatomy of dendritic spines using microinjection and confocal microscopy. Nature Protocols. 6:1391.

Ehrlich I, Klein M, Rumpel S, Malinow R. 2007. PSD-95 is required for activity-driven synapse stabilization. Proceedings of the National Academy of Sciences. 104:4176–4181.

El-Husseini AED, Schnell E, Chetkovich DM, Nicoll RA, Bredt DS. 2000. PSD-95 involvement in maturation of excitatory synapses. Science. 290:1364–1368.

Engert F, Bonhoeffer T. 1999. Dendritic spine changes associated with hippocampal long-term synaptic plasticity. Nature. 399:66–70.

Eroglu C, Barres BA. 2010. Regulation of synaptic connectivity by glia. Nature. 468:223– 231.

Fauth M, Tetzlaff C. 2016. Opposing effects of neuronal activity on structural plasticity. Frontiers in Neuroanatomy. 10:75.

Fossati G, Morini R, Corradini I, Antonucci F, Trepte P, Edry E, Sharma V, Papale A, Pozzi D, Defilippi P, et al. 2015. Reduced SNAP-25 increases PSD-95 mobility and impairs spine morphogenesis. Cell Death & Differentiation. 22:1425–1436.

Franklin KB, Paxinos G, et al. 2008. The mouse brain in stereotaxic coordinates. Academic Press.

Fremeau RTJ, Voglmaier S, Seal RP, Edwards RH. 2004. VGLUTs define subsets of excitatory neurons and suggest novel roles for glutamate. Trends in Neurosciences. 27:98–103.

Gainey MA, Feldman DE. 2017. Multiple shared mechanisms for homeostatic plasticity in rodent somatosensory and visual cortex. Philosophical Transactions of the Royal Society B: Biological Sciences. 372:20160157.

Gallinaro JV, Gasparovic N, Rotter S. 2020. Homeostatic structural plasticity leads to the formation of memory engrams through synaptic rewiring in recurrent networks. bioRxiv. .

Gallinaro JV, Rotter S. 2018. Associative properties of structural plasticity based on firing rate homeostasis in recurrent neuronal networks. Scientific Reports. 8:1–13.

Garrido-García A, Andrés-Pans B, Durán-Trío L, Díez-Guerra FJ. 2009. Activity-dependent translocation of neurogranin to neuronal nuclei. Biochemical Journal. 424:419–429.

Gipson CD, Olive MF. 2017. Structural and functional plasticity of dendritic spines–root or result of behavior? Genes, Brain and Behavior. 16:101–117.

Gray NW, Weimer RM, Bureau I, Svoboda K. 2006. Rapid redistribution of synaptic PSD-95 in the neocortex in vivo. PLOS Biology. 4:e370.

Han KS, Cooke SF, Xu W. 2017. Experience-dependent equilibration of ampar-mediated synaptic transmission during the critical period. Cell Reports. 18:892–904.

Haydon PG, Nedergaard M. 2015. How do astrocytes participate in neural plasticity? Cold Spring Harbor Perspectives in Biology. 7:a020438.

Hebb DO. 1949. The organization of behavior: a neuropsychological theory. J. Wiley; Chapman & Hall.

Hengen KB, Pacheco AT, McGregor JN, Van Hooser SD, Turrigiano GG. 2016. Neuronal firing rate homeostasis is inhibited by sleep and promoted by wake. Cell. 165:180–191.

Hol EM, Pekny M. 2015. Glial fibrillary acidic protein (GFAP) and the astrocyte intermediate filament system in diseases of the central nervous system. Current Opinion in Cell Biology. 32:121–130.

Holtmaat A, Svoboda K. 2009. Experience-dependent structural synaptic plasticity in the mammalian brain. Nature Reviews Neuroscience. 10:647.

Holtmaat A, Wilbrecht L, Knott GW, Welker E, Svoboda K. 2006. Experience-dependent and cell-type-specific spine growth in the neocortex. Nature. 441:979–983.

Huang KP, Huang FL, Shetty PK. 2011. Stimulation-mediated translocation of calmodulin and neurogranin from soma to dendrites of mouse hippocampal CA1 pyramidal neurons. Neuroscience. 178:1–12.

Humo M, Lu H, Yalcin I. 2019. The molecular neurobiology of chronic pain–induced depression. Cell and Tissue Research. :1–23.

Hwang H, Szucs MJ, Ding LJ, Allen A, Ren X, Haensgen H, Gao F, Rhim H, Andrade A, Pan JQ, et al. 2021. Neurogranin, encoded by the schizophrenia risk gene NRGN, bidirectionally modulates synaptic plasticity via calmodulin-dependent regulation of the neuronal phosphoproteome. Biological Psychiatry. 89:256–269.

Jo S, Yarishkin O, Hwang YJ, Chun YE, Park M, Woo DH, Bae JY, Kim T, Lee J, Chun H, et al. 2014. GABA from reactive astrocytes impairs memory in mouse models of alzheimer’s disease. Nature Medicine. 20:886.

Keck T, Mrsic-Flogel TD, Afonso MV, Eysel UT, Bonhoeffer T, Hübener M. 2008. Massive restructuring of neuronal circuits during functional reorganization of adult visual cortex. Nature Neuroscience. 11:1162–1167.

Keck T, Toyoizumi T, Chen L, Doiron B, Feldman DE, Fox K, Gerstner W, Haydon PG, Hübener M, Lee HK, et al. 2017. Integrating hebbian and homeostatic plasticity: the current state of the field and future research directions. Philosophical Transactions of the Royal Society B: Biological Sciences. 372:20160158.

Knott GW, Quairiaux C, Genoud C, Welker E. 2002. Formation of dendritic spines with gabaergic synapses induced by whisker stimulation in adult mice. Neuron. 34:265–273.

Koga K, Descalzi G, Chen T, Ko HG, Lu J, Li S, Son J, Kim T, Kwak C, Huganir RL, et al. 2015. Coexistence of two forms of LTP in ACC provides a synaptic mechanism for the interactions between anxiety and chronic pain. Neuron. 85:377–389.

Konur S, Rabinowitz D, Fenstermaker VL, Yuste R. 2003. Systematic regulation of spine sizes and densities in pyramidal neurons. Journal of Neurobiology. 56:95–112.

Lamprecht R, LeDoux J. 2004. Structural plasticity and memory. Nature Reviews Neuroscience. 5:45–54.

Lazar A, Pipa G, Triesch J. 2009. SORN: a self-organizing recurrent neural network. Frontiers in computational neuroscience. 3:23.

Lenz M, Galanis C, Kleidonas D, Fellenz M, Deller T, Vlachos A. 2019. Denervated mouse dentate granule cells adjust their excitatory but not inhibitory synapses following in vitro entorhinal cortex lesion. Experimental Neurology. 312:1–9.

Letellier M, Levet F, Thoumine O, Goda Y. 2019. Differential role of pre-and postsynaptic neurons in the activity-dependent control of synaptic strengths across dendrites. PLOS Biology. 17:e2006223.

Liddelow SA, Barres BA. 2017. Reactive astrocytes: production, function, and therapeutic potential. Immunity. 46:957–967.

Linssen C, Lepperød ME, Mitchell J, Pronold J, Eppler JM, Keup C, Peyser A, Kunkel S, Weidel P, Nodem Y, Terhorst D, Deepu R, Deger M, Hahne J, Sinha A, Antonietti A, Schmidt M, Paz L, Garrido J, Ippen T, Riquelme L, Serenko A, Kühn T, Kitayama I, Mørk H, Spreizer S, Jordan J, Krishnan J, Senden M, Hagen E, Shusharin A, Vennemo SB, Rodarie D, Morrison A, Graber S, Schuecker J, Diaz S, Zajzon B, Plesser HE. 2018. NEST 2.16.0. doi:10.5281/zenodo.1400175.

Litwin-Kumar A, Doiron B. 2014. Formation and maintenance of neuronal assemblies through synaptic plasticity. Nature Communications. 5:1–12.

Liu X, Ramirez S, Pang PT, Puryear CB, Govindarajan A, Deisseroth K, Tonegawa S. 2012. Optogenetic stimulation of a hippocampal engram activates fear memory recall. Nature. 484:381–385.

Lowel S, Singer W. 1992. Selection of intrinsic horizontal connections in the visual cortex by correlated neuronal activity. Science. 255:209–212.

Lu H, Gallinaro JV, Rotter S. 2019. Network remodeling induced by transcranial brain stimulation: A computational model of tdcs-triggered cell assembly formation. Network Neuroscience. 3:924–943.

Lucassen PJ, Pruessner J, Sousa N, Almeida OF, Van Dam AM, Rajkowska G, Swaab DF, Cźeh B. 2014. Neuropathology of stress. Acta Neuropathologica. 127:109–135.

Lynch GS, Dunwiddie T, Gribkoff V. 1977. Heterosynaptic depression: a postsynaptic correlate of long-term potentiation. Nature. 266:737–739.

Ma Z, Turrigiano GG, Wessel R, Hengen KB. 2019. Cortical circuit dynamics are homeostatically tuned to criticality in vivo. Neuron. 104:655–664.

Malenka RC, Bear MF. 2004. LTP and LTD: an embarrassment of riches. Neuron. 44:5–21.

Markram H, Lübke J, Frotscher M, Sakmann B. 1997. Regulation of synaptic efficacy by coincidence of postsynaptic APs and EPSPs. Science. 275:213–215.

Matsuzaki M, Honkura N, Ellis-Davies GC, Kasai H. 2004. Structural basis of long-term potentiation in single dendritic spines. Nature. 429:761–766.

Miller KD, MacKay DJ. 1994. The role of constraints in Hebbian learning. Neural Computation. 6:100–126.

Murphy TH, Corbett D. 2009. Plasticity during stroke recovery: from synapse to behaviour. Nature Reviews Neuroscience. 10:861–872.

Nagel G, Szellas T, Huhn W, Kateriya S, Adeishvili N, Berthold P, Ollig D, Hegemann P, Bamberg E. 2003. Channelrhodopsin-2, a directly light-gated cation-selective membrane channel. Proceedings of the National Academy of Sciences. 100:13940–13945.

Nikolic K, Jarvis S, Grossman N, Schultz S. 2013. Computational models of optogenetic tools for controlling neural circuits with light. In: 2013 35th Annual International Conference of the IEEE Engineering in Medicine and Biology Society (EMBC). IEEE. p. 5934–5937.

Noguchi J, Nagaoka A, Hayama T, Ucar H, Yagishita S, Takahashi N, Kasai H. 2019. Bidirectional in vivo structural dendritic spine plasticity revealed by two-photon glutamate uncaging in the mouse neocortex. Scientific Reports. 9:1–8.

Nollet M, Guisquet AML, Belzung C. 2013. Models of depression: unpredictable chronic mild stress in mice. Current Protocols in Pharmacology. 61:5–65.

Ohsawa K, Imai Y, Sasaki Y, Kohsaka S. 2004. Microglia/macrophage-specific protein Iba1 binds to fimbrin and enhances its actin-bundling activity. Journal of Neurochemistry. 88:844–856.

Pacheco AT, Tilden EI, Grutzner SM, Lane BJ, Wu Y, Hengen KB, Gjorgjieva J, Turrigiano GG. 2019. Rapid and active stabilization of visual cortical firing rates across light–dark transitions. Proceedings of the National Academy of Sciences. 116:18068– 18077.

Pak DT, Sheng M. 2003. Targeted protein degradation and synapse remodeling by an inducible protein kinase. Science. 302:1368–1373.

Pfeiffer T, Poll S, Bancelin S, Angibaud J, Inavalli VK, Keppler K, Mittag M, Fuhrmann M, Nägerl UV. 2018. Chronic 2P-STED imaging reveals high turnover of dendritic spines in the hippocampus in vivo. eLife. 7:e34700

R Core Team. 2019. R: A Language and Environment for Statistical Computing. R Foundation for Statistical Computing. Vienna, Austria.

Ramirez S, Liu X, Lin PA, Suh J, Pignatelli M, Redondo RL, Ryan TJ, Tonegawa S. 2013. Creating a false memory in the hippocampus. Science. 341:387–391.

Ryan TJ, Roy DS, Pignatelli M, Arons A, Tonegawa S. 2015. Engram cells retain memory under retrograde amnesia. Science. 348:1007–1013.

Samuels BA, Hen R. 2011. Novelty-suppressed feeding in the mouse. In: Mood and anxiety related phenotypes in mice. Springer. p. 107–121.

Sanderson TM, Georgiou J, Collingridge GL. 2020. Illuminating relationships between the pre-and post-synapse. Frontiers in Neural Circuits. 14.

Schousboe A, Waagepetersen HS. 2005. Role of astrocytes in glutamate homeostasis: implications for excitotoxicity. Neurotoxicity Research. 8:221–225.

Sejnowski TJ. 1977. Storing covariance with nonlinearly interacting neurons. Journal of Mathematical Biology. 4:303–321.

Sellmeijer J, Mathis V, Hugel S, Li XH, Song Q, Chen QY, Barthas F, Lutz PE, Karatas M, Luthi A, et al. 2018. Hyperactivity of anterior cingulate cortex areas 24a/24b drives chronic pain-induced anxiodepressive-like consequences. Journal of Neuroscience. 38:3102–3115.

Sholl DA. 1953. Dendritic organization in the neurons of the visual and motor cortices of the cat. Journal of Anatomy. 87:387.

Shu Y, Hasenstaub A, McCormick DA. 2003. Turning on and off recurrent balanced cortical activity. Nature. 423:288–293.

Singec I, Knoth R, Ditter M, Volk B, Frotscher M. 2004. Neurogranin is expressed by principal cells but not interneurons in the rodent and monkey neocortex and hippocampus. Journal of Comparative Neurology. 479:30–42

Steinmetz CC, Turrigiano GG. 2010. Tumor necrosis factor-*α* signaling maintains the ability of cortical synapses to express synaptic scaling. Journal of Neuroscience. 30:14685– 14690.

Stellwagen D, Malenka RC. 2006. Synaptic scaling mediated by glial tnf-*α*. Nature. 440:1054–1059.

Svirsky S, Henchir J, Li Y, Ma X, Carlson S, Dixon CE. 2020. Neurogranin protein expression is reduced after controlled cortical impact in rats. Journal of Neurotrauma. 37:939–949.

Toyoizumi T, Kaneko M, Stryker MP, Miller KD. 2014. Modeling the dynamic interaction of Hebbian and homeostatic plasticity. Neuron. 84:497–510.

Trachtenberg JT, Chen BE, Knott GW, Feng G, Sanes JR, Welker E, Svoboda K. 2002. Long-term in vivo imaging of experience-dependent synaptic plasticity in adult cortex. Nature. 420:788–794.

Turrigiano G. 2012. Homeostatic synaptic plasticity: local and global mechanisms for stabilizing neuronal function. Cold Spring Harbor Perspectives in Biology. 4:a005736.

Turrigiano GG. 2017. The dialectic of hebb and homeostasis. Philosophical Transactions of the Royal Society B: Biological Sciences. 372:20160258.

Turrigiano GG, Leslie KR, Desai NS, Rutherford LC, Nelson SB. 1998. Activity-dependent scaling of quantal amplitude in neocortical neurons. Nature. 391:892–896.

Van Ooyen A. 2011. Using theoretical models to analyse neural development. Nature Reviews Neuroscience. 12:311–326.

Van Vreeswijk C, Sompolinsky H. 1996. Chaos in neuronal networks with balanced excitatory and inhibitory activity. Science. 274:1724–1726.

Vlachos A, Müller-Dahlhaus F, Rosskopp J, Lenz M, Ziemann U, Deller T. 2012a. Repetitive magnetic stimulation induces functional and structural plasticity of excitatory postsynapses in mouse organotypic hippocampal slice cultures. Journal of Neuroscience. 32:17514–17523.

Vlachos A, Orth CB, Schneider G, Deller T. 2012b. Time-lapse imaging of granule cells in mouse entorhino-hippocampal slice cultures reveals changes in spine stability after entorhinal denervation. Journal of Comparative Neurology. 520:1891–1902.

Vogels TP, Froemke RC, Doyon N, Gilson M, Haas JS, Liu R, Maffei A, Miller P, Wierenga C, Woodin MA, et al. 2013. Inhibitory synaptic plasticity: spike timing-dependence and putative network function. Frontiers in Neural Circuits. 7:119.

Vogels TP, Sprekeler H, Zenke F, Clopath C, Gerstner W. 2011. Inhibitory plasticity balances excitation and inhibition in sensory pathways and memory networks. Science. 334:1569–1573.

Weinhard L, di Bartolomei G, Bolasco G, Machado P, Schieber NL, Neniskyte U, Exiga M, Vadisiute A, Raggioli A, Schertel A, et al. 2018. Microglia remodel synapses by presynaptic trogocytosis and spine head filopodia induction. Nature Communications. 9:1228.

Williams JC, Xu J, Lu Z, Klimas A, Chen X, Ambrosi CM, Cohen IS, Entcheva E. 2013. Computational optogenetics: empirically-derived voltage-and light-sensitive channelrhodopsin-2 model. PLOS Computational Biology. 9.

Wilson NR, Kang J, Hueske EV, Leung T, Varoqui H, Murnick JG, Erickson JD, Liu G. 2005. Presynaptic regulation of quantal size by the vesicular glutamate transporter vglut1. Journal of Neuroscience. 25:6221–6234.

Xia Z, Storm DR. 2005. The role of calmodulin as a signal integrator for synaptic plasticity. Nature Reviews Neuroscience. 6:267–276.

Yizhar O, Fenno LE, Davidson TJ, Mogri M, Deisseroth K. 2011. Optogenetics in neural systems. Neuron. 71:9–34.

Yusifov R, Tippmann A, Staiger JF, Schlüter OM, Löwel S. 2021. Spine dynamics of PSD-95-deficient neurons in the visual cortex link silent synapses to structural cortical plasticity. Proceedings of the National Academy of Sciences. 118.

Zamani A, Sakuragi S, Ishizuka T, Yawo H. 2017. Kinetic characteristics of chimeric channelrhodopsins implicate the molecular identity involved in desensitization. Biophysics and Physicobiology. 14:13–22.

Zamanian JL, Xu L, Foo LC, Nouri N, Zhou L, Giffard RG, Barres BA. 2012. Genomic analysis of reactive astrogliosis. Journal of Neuroscience. 32:6391–6410.

Zenke F, Gerstner W. 2017. Hebbian plasticity requires compensatory processes on multiple timescales. Philosophical Transactions of the Royal Society B: Biological Sciences. 372:20160259.

Zhang W, Vazquez L, Apperson M, Kennedy MB. 1999. Citron binds to PSD-95 at glutamatergic synapses on inhibitory neurons in the hippocampus. Journal of Neuroscience. 19:96–108.

Zhong L, Gerges NZ. 2012. Neurogranin targets calmodulin and lowers the threshold for the induction of long-term potentiation. PLOS ONE. 7.

Zhou Q, Homma KJ, Poo Mm. 2004. Shrinkage of dendritic spines associated with long-term depression of hippocampal synapses. Neuron. 44:749–757.

Zuur A, Ieno EN, Walker N, Saveliev AA, Smith GM. 2009. Mixed effects models and extensions in ecology with R. Springer Science & Business Media.

