## Supplementary Materials for "Time course of homeostatic structural plasticity in response to optogenetic stimulation in mouse anterior cingulate cortex"

### Virus injection

To compare the effects of optogenetic stimulation between the transgenic animals and viral injected ones, we also transfected C57BL/6J mice with virus pAAV-CaMKIIa-hChR2(H134R)-EYFP (Addgene plasmid #26969; `http://n2t.net/addgene:26969`; RRID: Addgene_26969). We injected the same virus bilaterally into the ACC (0.5 $\mu$L per side, with a speed of 0.05 $\mu$L per 30 seconds) in both stimulated and sham mice. 15 min after the injection, We removed the Hamilton syringe from the brain. The same stimulation protocol and behavioral tests were used for both transfected mice and transgenic mice.

### Tissue harvesting

For immunohistochemical staining, the perfusion protocol contains 2 min wash with cold 1× phosphate buffer (PB, 0.1 M, pH 7.4) solution and 18 min perfusion with 4% paraformaldehyde (PFA) in 1× PB (with the pump speed at 10 mL/min). Then brains were extracted and kept in 4% PFA in 1× PB overnight for post-fixation. Brains were later transferred and stored in 1× PBS with 0.02% sodium azide at 4 °C for further experiments.

For microinjection experiments, we used a fast perfusion method to protect the fine dendritic structure (Dumitriu et al. 2011). We washed the blood with 1% PFA in 1× PB for 2 min, and perfused animals for another 12 min with 4% PFA in 1× PB with 0.125% glutaraldehyde. The pump speed was 5 mL/min. Then we extracted the brains and put them in 4% PFA in 1× PB with 0.125% glutaraldehyde for another 6 hrs for post-fixation. All brains were later transferred and stored in 1× PBS with 0.02% sodium azide at 4 °C for further experiments.

### Linear mixed models

In the current study, we aimed to analyze the difference between the sham and stimulated mice, in terms of the expression of markers and neural morphology. Besides, we wanted to specify if the difference between the stimulated and sham groups showed various patterns on the ipsi- and contralateral hemispheres. In principle, these analyses are analyses of variance (ANOVA) or two-way ANOVA, depending on the number of explanatory variables. Applying ANOVA, however, demands the satisfaction of certain assumptions. The relationship between the explanatory variables and the response variable should be linear. The independence of observations ought to be ensured. Apart from that, the variance of observations should be normally distributed and remain homogeneous for different groups. In the current study, however, each mouse either contributed several sections for each distance group or provided several neurons, which violated the independence assumption. The distribution of data for each mouse or each group is not necessarily homogeneous. In such conditions, initial trials with Mann Whitney U test or Kruskal Wallis test returned appropriate highly significant results and risked our study for achieving inappropriate conclusions. So we used a new method to deal with the heterogeneity and dependence of our data points, the linear mixed model (LMM).

**IHC quantification and LMM**

In an LMM model, we specified the effects of fixed and random components. Fixed effects refer to the effects explained by the explanatory factors, while random effects refer to the nestedness of data. With visual inspection, we used being stimulated or not (sham/stimulated) and stimulation side (ipsi/contra) and their interaction effect as the fixed term to explain their difference. Animal ID or section ID and distance to the optic fiber are the random term. With this model, we can approximate the effects of explanatory variables as much as possible with all the data points collected from multiple mice and meanwhile control the discrepancy between mice.

In our study, we used LMM to analyze the results of two experiments, the immunohistochemical staining and microinjection, with different fixed effects and random effects structures. Here we showed the analysis of immunohistochemical staining results as a detailed example. The analysis of morphological data was similar and we will skip the details.

We quantified the protein signal intensity of both hemispheres of ACC sections from five distance groups in both sham and stimulated mice groups. We **hypothesized** that the overall signal intensity is different in the stimulated mice from the sham group. Moreover, the ipsilateral hemisphere of the stimulated mice should be different from the contralateral hemisphere, too, since the ipsilateral side is closer to the optic fiber and receives more stimulation. The contribution of stimulation side (ipsi/contra) to the signal intensity of the sham mice, in theory, would not be as robust as in the stimulated group. Thus, we proposed two explanatory variables and their interaction effects as the master structure of the fixed effects. As for random effects, we used mice and distance group as the group factor for sections. As showed in Equation S.1,

$$\text{Signal\_intensity} \sim \text{Sham} * \text{Ipsi} + (1|\text{AnimalID}/\text{Distance2}) + \epsilon, \tag{S.1}$$

*Signal_intensity* is the response variable. The first part follow the symbol $\sim$ is the fixed effects structure where *Sham* and *Ipsi* are two nominal explanatory variables and $*$ denoted the interaction effects between them. (1|AnimalID/Distabce2) is the random effects structure, grouping sections per mouse and per distance group. $\epsilon$ is the residual.

After fitted the model, we plotted the fitted values' residuals against each explanatory variable to inspect if the model fits well visually. The significance of fixed effects was tested by profiling the confidence interval of each estimated parameter. $p < 0.05$, $p < 0.01$, and $p < 0.001$ means 95% CI, 99% CI, or 99.9% CI of the estimated coefficient does not cross zero.

#### Counting data and GLMM

In the case of counting data, such as IBA1+ and c-Fos+ cell counting, Poisson distribution of data points was assumed, so LMM was no longer suitable. We used the generalized linear mixed model (GLMM) to fit the counting data. The same process, as described above, was used to estimate the parameters. In the last step, in addition to visual inspection, we also applied the over-dispersion test to validate our model. In the case of over-dispersion

(> 2.0) or underdispersion (< 0.6), we adjusted the model back and forth. Significance notification for the coefficients estimated by GLMM models remained the same as LMM models.

#### Morphological analysis and LMM

Different from immunohistochemical staining data, we did not specify distance information for each neuron. All the neurons were selected from sections within ±0.4 mm anterior-posterior (AP) away from the optic fiber and treated as homogeneous. So regarding the random effects structure, we used two types depending on the response variable. If quantifications were taken per neuron, such as soma size and Sholl intersections, we used mice as the group factor (model1). If quantifications were taken per dendrite segment of individual neurons for each mouse, such as the dendritic length, average dendritic diameter, and spine density, we used a two-layer group factor with both neurons and mice (model2).

```
model1 <- lmer(Soma _size~Sham*Ipsi + (1|AnimalID),REML=TRUE,data=time24h)
model2 <- lmer(Spine_density~Sham*Ipsi+(1|AnimalID:NeuronID),REML=TRUE,data=time24h)
```

#### Comparing LMM with other tests

Here in this section, we showed an example dataset and compared the results of different analysis. Supplementary Figure 4-1 showed the distribution of stubby spine head volume from the apical dendrites for each mouse sacrificed at 24 h post-stimulation. Visually we saw each mouse contributed several spines from various segments. If we ignored the independence assumption and pooled all the data points, the difference between sham and the stimulated mice was significant ($p = 0.0002$, the Kolmogorov-Smirnov test; $p = 2.8 * 10^{-7}$, Mann Whitney U test). If we specified the random effects and grouped the data points by the dendrite segment and by mouse, the sham and the stimulated mice were no longer significantly different from each other ($p > 0.05$, LMM) as presented in Figure 4G (upper panel). So breaking the independence assumption could result in inappropriate conclusions.

### Neuron model

The membrane potential dynamics of linear integrate-and-fire (LIF) neuron model is explained by Equation S.2,

$$\tau_m \frac{\mathrm{d}}{\mathrm{dt}} V_i(t) = -V_i(t) + \tau_m \sum_j J_{ij} S_j(t-d) + J_{\mathrm{opto}} S_{\mathrm{opto}}(t-d), \qquad \text{(S.2)}$$

where $V_i(t)$ is the membrane potential of neuron $i$, with a resting value at $0\,\mathrm{mV}$. $\tau_m$ is the membrane time constant. $S_j(t) = \sum_k \delta(t - t_j^k)$ represents the spike train generated by neuron $j$, where $t_j^k$ represents the individual spike times, and $d$ is the synaptic transmission delay. The amplitude of the postsynaptic potential that induced in neuron $i$ upon the arrival of a spike from neuron $j$ is governed by matrix $J_{ij}$. $S_{\mathrm{opto}}(t-d)$ represents the depolarizing inputs from optotgenetic stimulation to a subset of excitatory neurons with synaptic weight $J_{\mathrm{opto}}$. When the membrane potential $V_i(t)$ reaches the threshold, $V_{\mathrm{th}}$, an action potential is emitted and the membrane potential is reset to $V_{\mathrm{reset}} = 10\,\mathrm{mV}$. All parameters of neuron model are summarized in Supplementary Table 4.

### Network model

ACC was modeled as an inhibition-dominated recurrent network (Brunel 2000), including 10 000 excitatory and 2 500 inhibitory neurons. The excitatory synapses onto inhibitory neurons (E-I) and all inhibitory connections (I-I and I-E) were static with a fixed weight. All excitatory synapses are at $J_E = 0.1\,\mathrm{mV}$ , while all inhibitory synapses are at $J_I = -0.8\,\mathrm{mV}$. These connections were beforehand randomly formed with a 10% connection probability. On the other hand, the excitatory-to-excitatory (E-E) connections were grown from zero based on the rule of homeostatic structural plasticity (Diaz-Pier et al. 2016; Gallinaro and Rotter 2018). Each neuron in the network connected to a Poissonian external input at a rate of $r_{\mathrm{ext}} = 30\,\mathrm{kHz}$. E-E connections formed during the growth period, and the network automatically entered an asynchronous-irregular state (Brunel 2000) when the equilibrium state were reached. All network parameters are listed in Supplementary Table 5.

### Homeostatic structural plasticity model

Homeostatic structural plasticity rule governed the growth of presynaptic elements (boutons) and postsynaptic elements (spines) via calcium concentration, $C(t) = [\mathrm{Ca}^{2+}]$. As showed in Equation S.3, whenever the neuron emits a spike, the intracellular calcium concentration experiences an influx by the amount $\beta_{\mathrm{Ca}}$. Between spikes, the calcium concentration decays exponentially with time constant $\tau_{\mathrm{Ca}}$.

$$\frac{\mathrm{d}}{\mathrm{dt}}C(t) = -\frac{1}{\tau_{\mathrm{Ca}}}C(t) + \beta_{\mathrm{Ca}}S(t). \tag{S.3}$$

We applied a linear growth rule to both pre- and postsynaptic elements (Equation S.4). Each neuron tends to maintain its activity as well as calcium concentration at a homeostatic level, called *set-point*. When the firing rate (or calcium concentration) falls below its set-point, the neuron will grow new synaptic elements and form functional synapses. When the firing rate (or calcium concentration) goes above the set-point, the neuron will randoly break synapses and retract synaptic elements. The respective counter-parts of deleted elements are added to the pool of free synaptic elements. New synapses can form only if free synaptic elements are available. Pairs of neurons can form multiple synapses between them, and each individual functional synapse has the same weight $J_E = 0.1\,\mathrm{mV}$.

$$\frac{\mathrm{d}}{\mathrm{dt}}z(t) = \nu\big[1 - \frac{1}{\epsilon}C(t)\big], \tag{S.4}$$

where $z(t)$ is the total number of pre- or postsynaptic elements a neuron has, $\nu$ is the growth rate, and $\epsilon$ is the target level of calcium concentration. All the parameters about the structural plasticity rule are summarized in Supplementary Table 6.

### Measurements for the numerical experiments

#### Firing rate

The firing rate of a neuron was calculated from its spike count, in a 5 s activity recording. The mean firing rate of a population was taken to be the arithmetic mean of firing rates across neurons in the population.

### Connectivity

Let $(A_{ij})$ be the $n \times n$ connectivity matrix of a network with $n$ neurons. Its columns correspond to the axons, its rows correspond to the dendrites of the neurons involved. The specific entry $A_{ij}$ of this matrix represents the total number of synapses from the presynaptic neuron $j$ to the postsynaptic neuron $i$. The mean connectivity of this network is then given by $\Gamma(t) = \frac{1}{n^2} \sum_{ij} A_{ij}$, where $t$ is the observing time point.

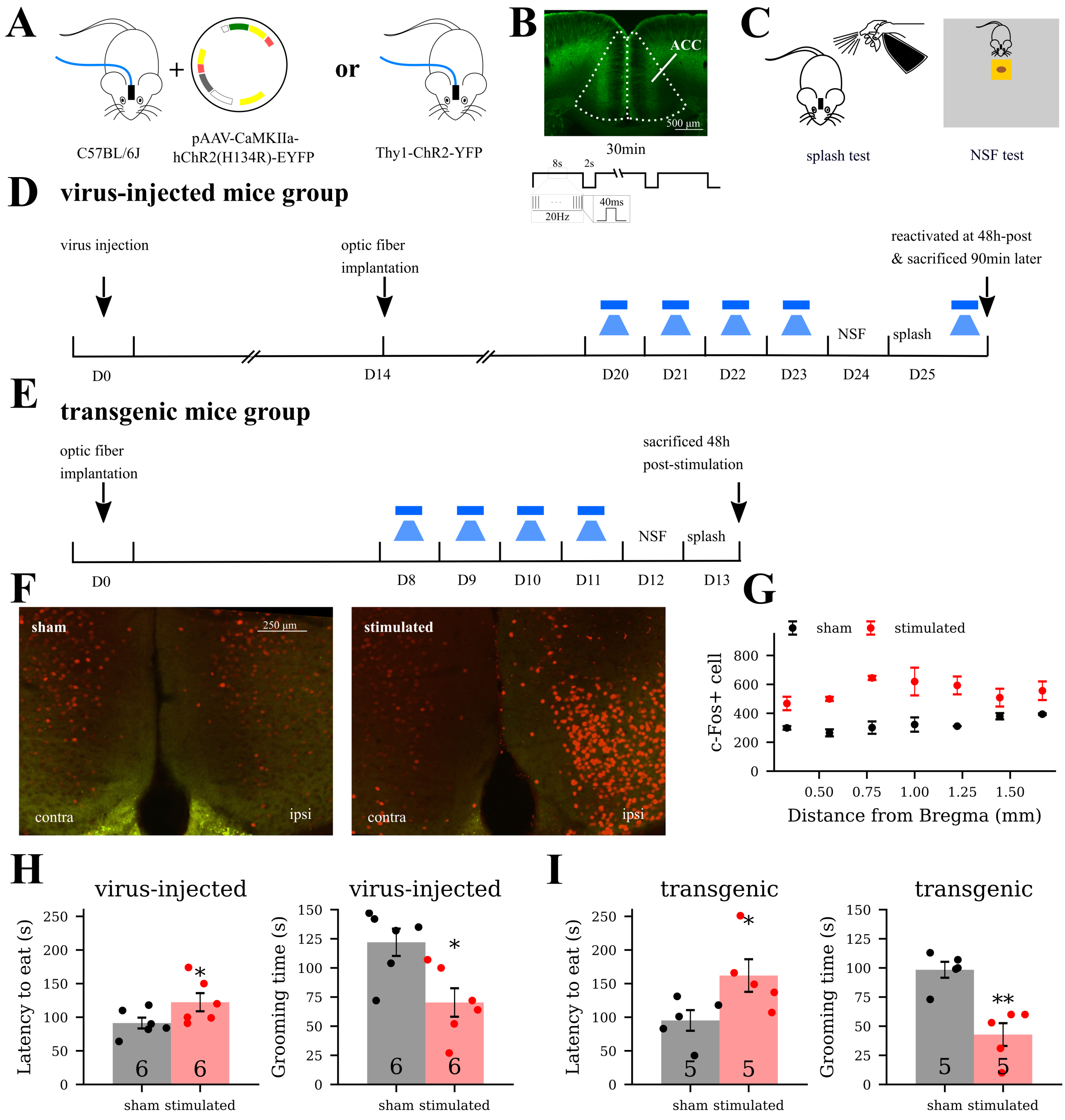


**Supplementary Figure 1-1:** No differences were observed between viral transfection and transgenic mice after the optogenetic stimulation at the behavioral and c-Fos expression level. **A** C57BL/6J mice bilaterally infected with virus vector and expressed hChR2 or Thy1-ChR2-YFP mice were used. **B-E** Both mice models received the same optogenetic stimulation protocol in ACC and behavior tests. **D** Virus-injected mice received activation again on the same day after splash test and sacrificed 1.5 h later to measure c-Fos expression. **F-G** c-Fos expression was elevated in the stimulated mice (99.5% CI = $[0.287, 0.687]$, GLMM). The c-Fos data was presented again in Figure 2G, left panel. **H-I** Mice in both approaches showed depressive-like behavior at 24 h and 48 h post-stimulation ($p = 0.012$ and $p = 0.032$ for the virus-injected group, $p = 0.018$ and $p = 0.0059$ for the transgenic group, Mann Whitney U test). The behavioral data from the transgenic mice (panel **I**) was presented again in Figure 1E-F.

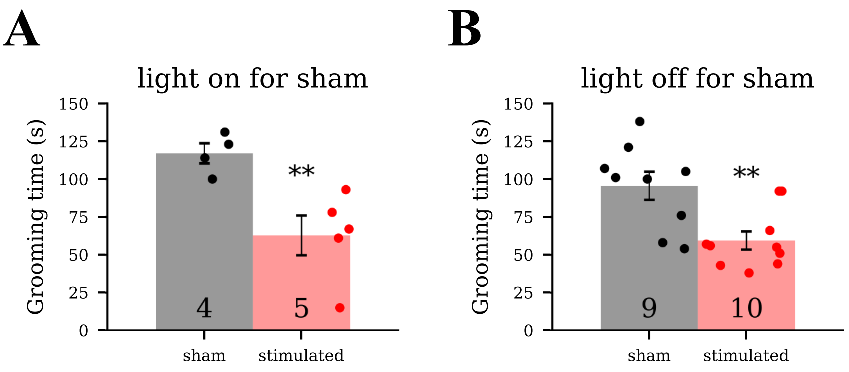


**Supplementary Figure 1-2:** Light did not trigger changes in behavior. **A** When Thy1-ChR2-YFP mice were used in both sham and stimulated groups, we kept the light off for sham mice during stimulation. Splash test showed stimulated mice presented depressive-like behavior after four days of stimulation ($p = 0.003$, Mann Whitney U test). **B** When Thy1-ChR2-YFP mice were used in the stimulated group while gene matched wild-type mice were used for sham, we kept the light on for the sham group during stimulation. Significant difference was observed in the splash test after four days of stimulation ($p = 0.009$, Mann Whitney U test). To keep the mice gene background consistent, we used Thy1-ChR2-YFP mice throughout our study and kept light off for the sham group.

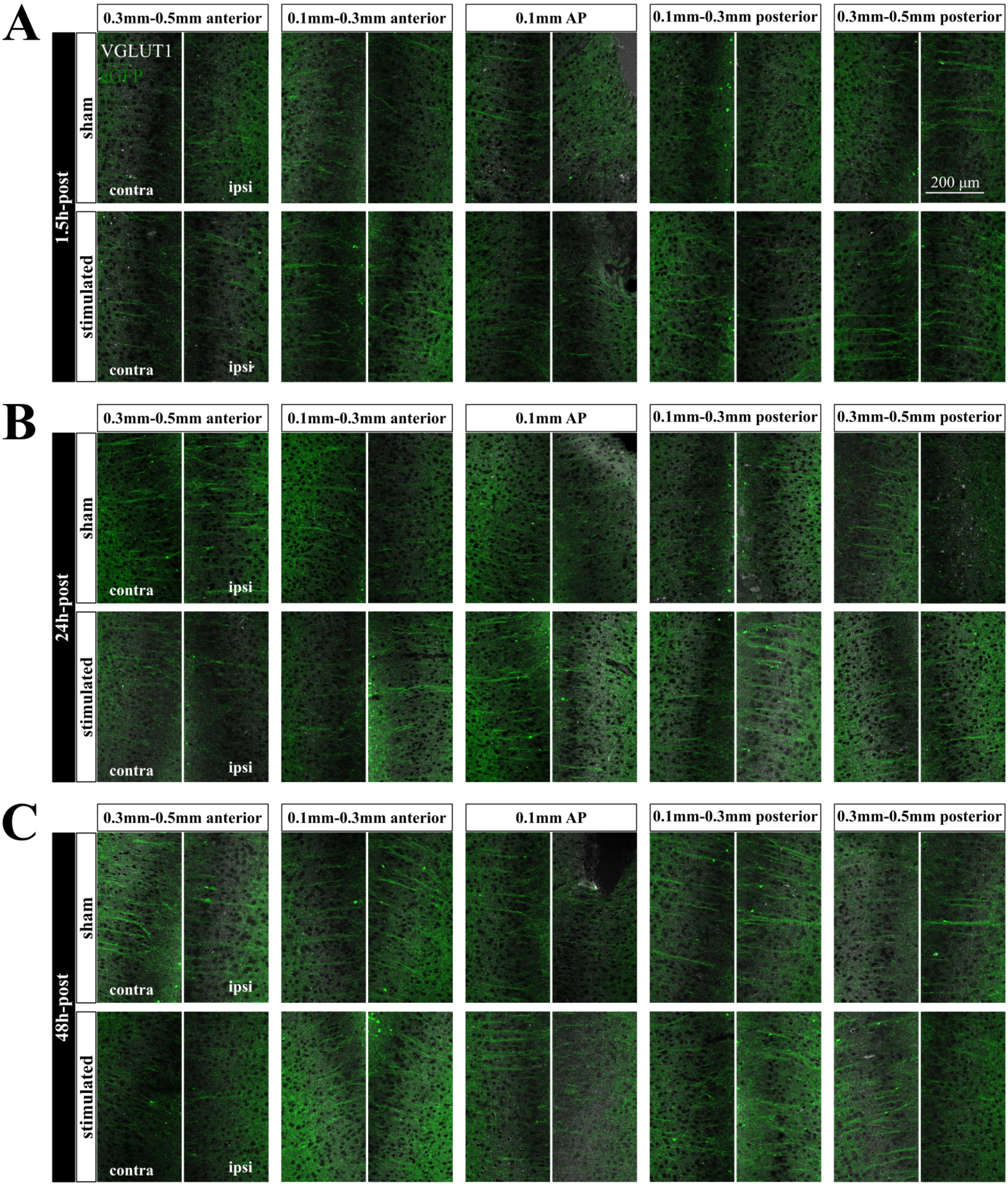


**Supplementary Figure 2-1:** Representative images of VGLUT1 staining at different distance to the optic fiber at 1.5 h, 24 h, and 48 h post-stimulation. High magnification images of the ipsilateral hemispheres in the middle panel were presented in Figure 2A.

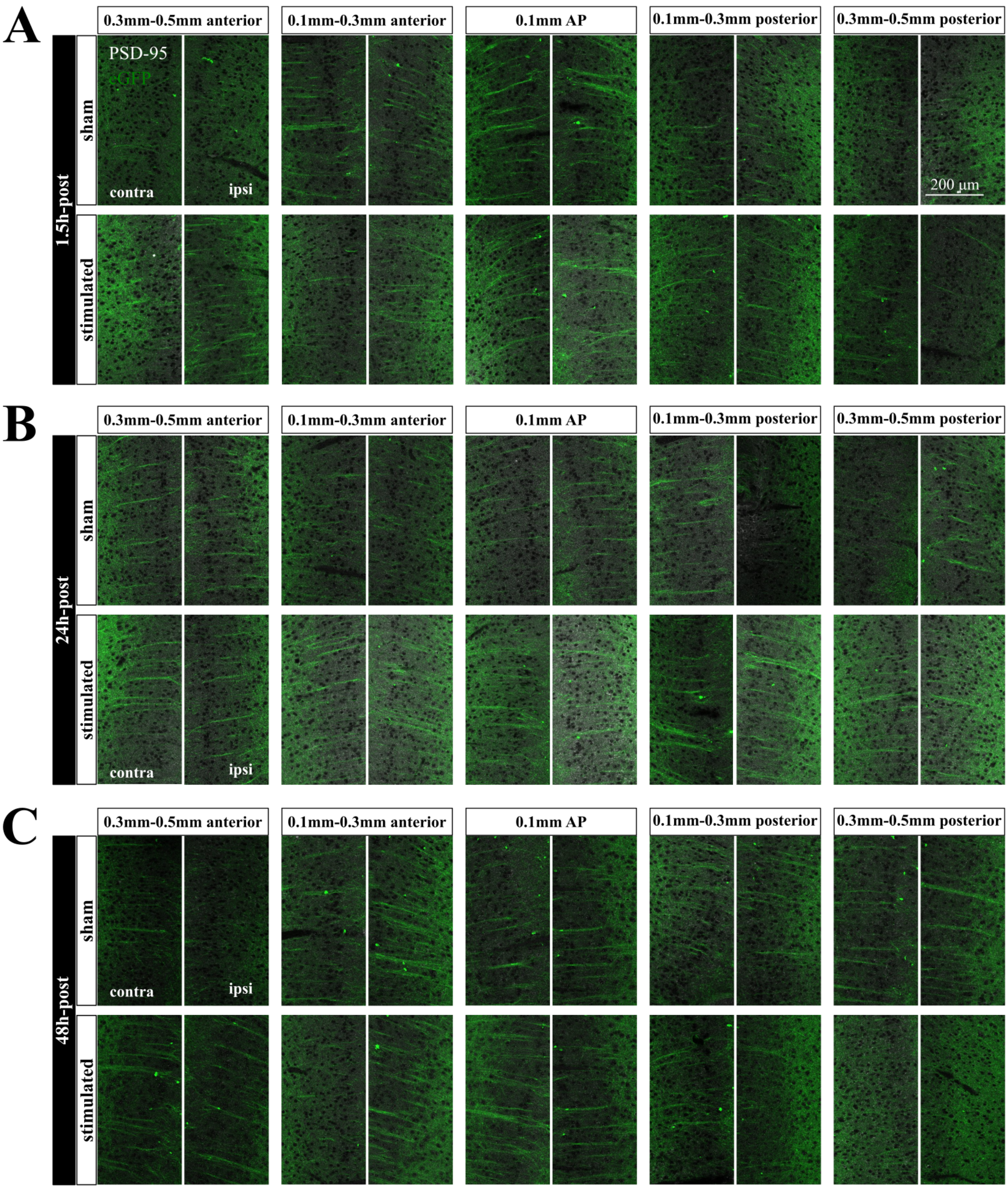


**Supplementary Figure 2-2:** Representative images of PSD-95 staining at different distance to the optic fiber at 1.5 h, 24 h, and 48 h post-stimulation. High magnification images of the ipsilateral hemispheres in the middle panel were presented in Figure 2C.

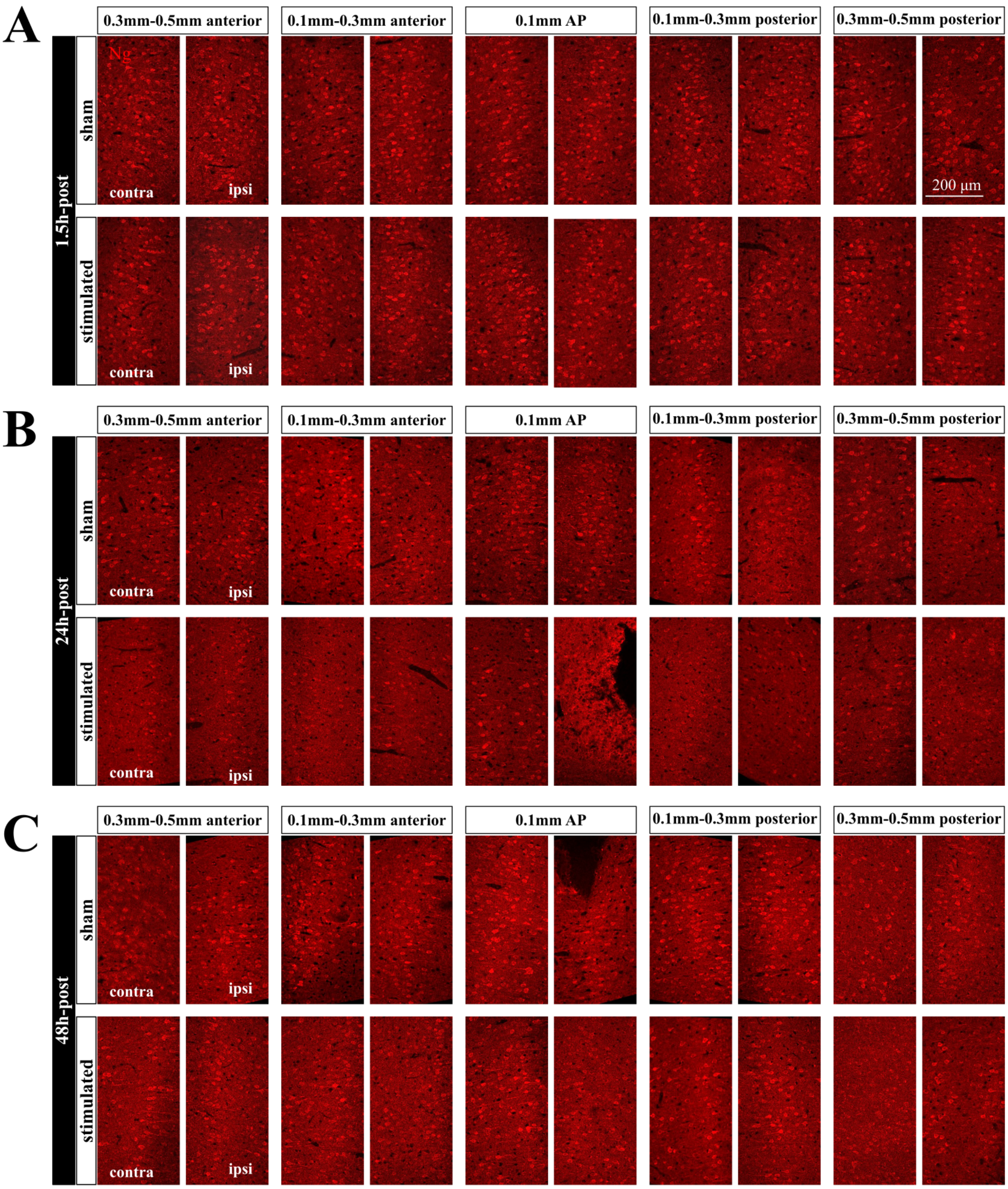


**Supplementary Figure 2-3:** Representative images of neurogranin staining at different distance to the optic fiber at 1.5 h, 24 h, and 48 h post-stimulation. High magnification images of the ipsilateral hemispheres in the middle panel were presented in Figure 2E.

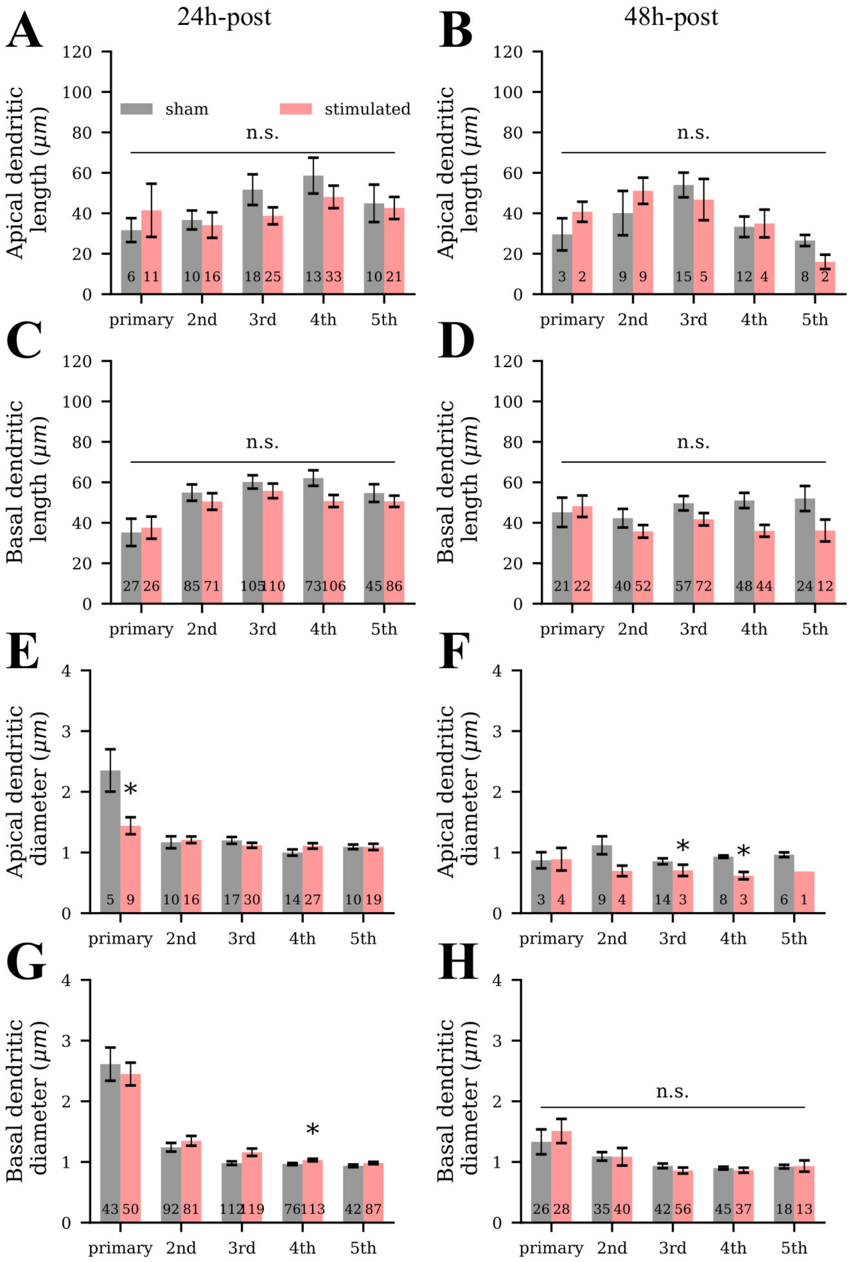


**Supplementary Figure 3-1:** Dendritic length and average dendritic diameter were not drastically altered at 24 h and 48 h post-stimulation. **A-D** Length of apical and basal dendrites were not altered by stimulation at 24 h and 48 h. **E-H** At 24 h, the average diameter of primary apical dendrites were reduced while the fourth level basal dendrites were increased by stimulation ($p < 0.05$ and $p < 0.05$). At 48 h, only the average diameter of third and fourth level apical dendrites were reduced by stimulation ($p < 0.05$ and $p < 0.05$). LMM was used for statistical analysis.

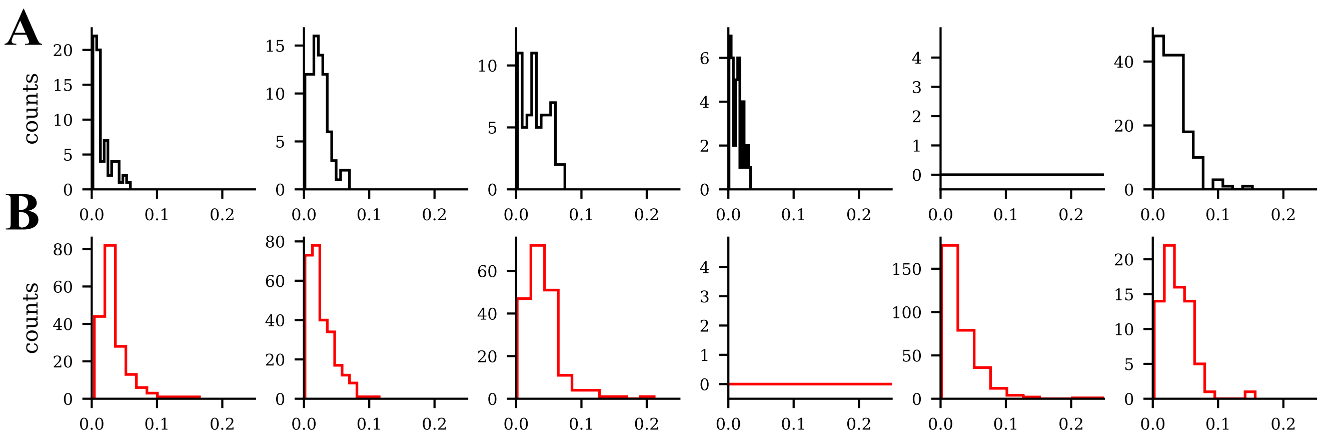


**Supplementary Figure 4-1:** The distribution of spine head volumes of stubby spins in the apical dendrites from each mouse at 24 h post-stimulation. The black histograms represent six sham mice respectively, while the red histograms represent stimulated mice. The fifth sham mouse and the fourth stimulated mouse did not contribute apical dendrite segments.

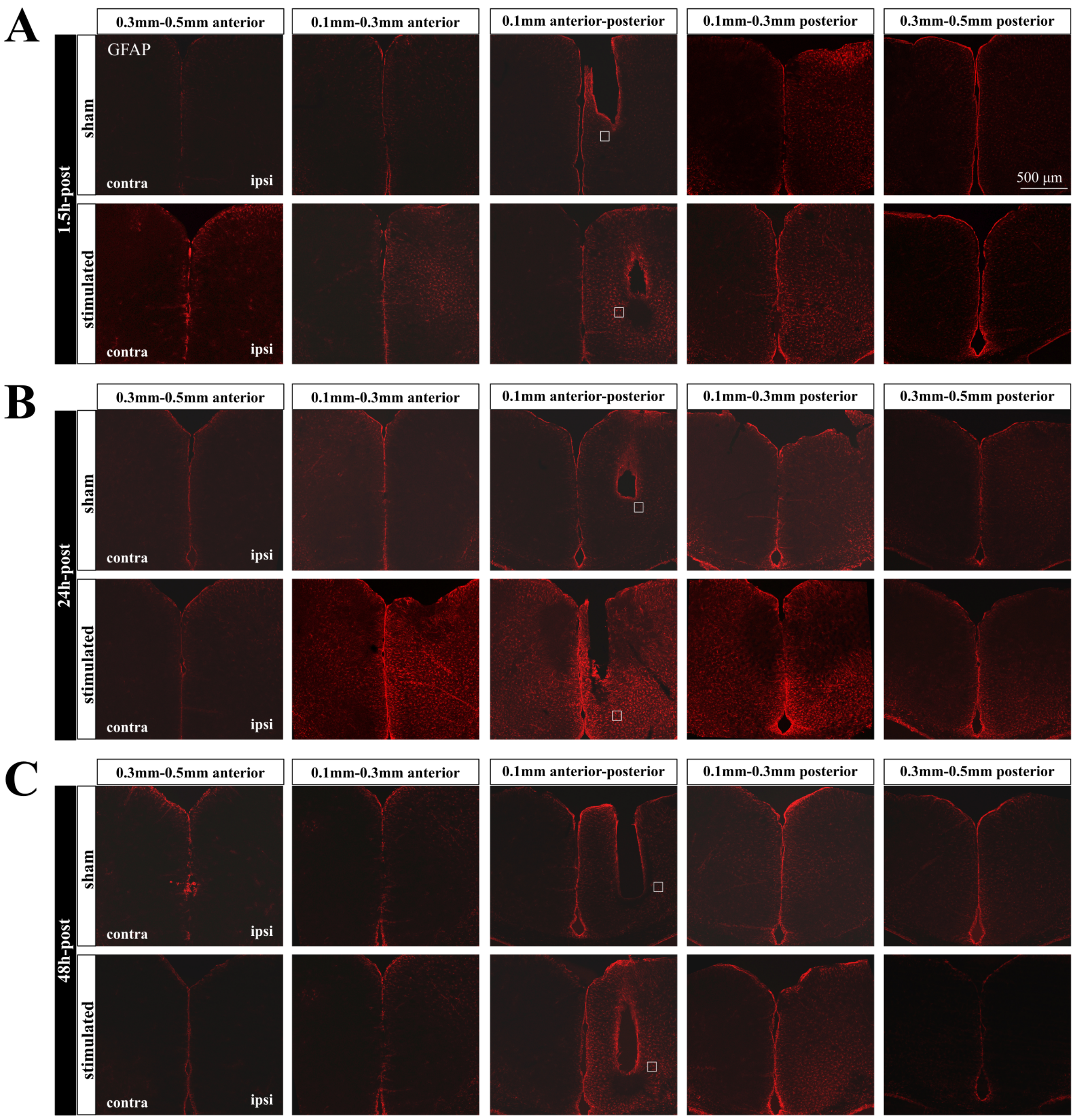


**Supplementary Figure 5-1:** Representative images of GFAP staining at different distance to the optic fiber at 1.5 h, 24 h, and 48 h post-stimulation. High magnification images of the selected regions (white squares) in the middle panel were presented in Figure 5A.

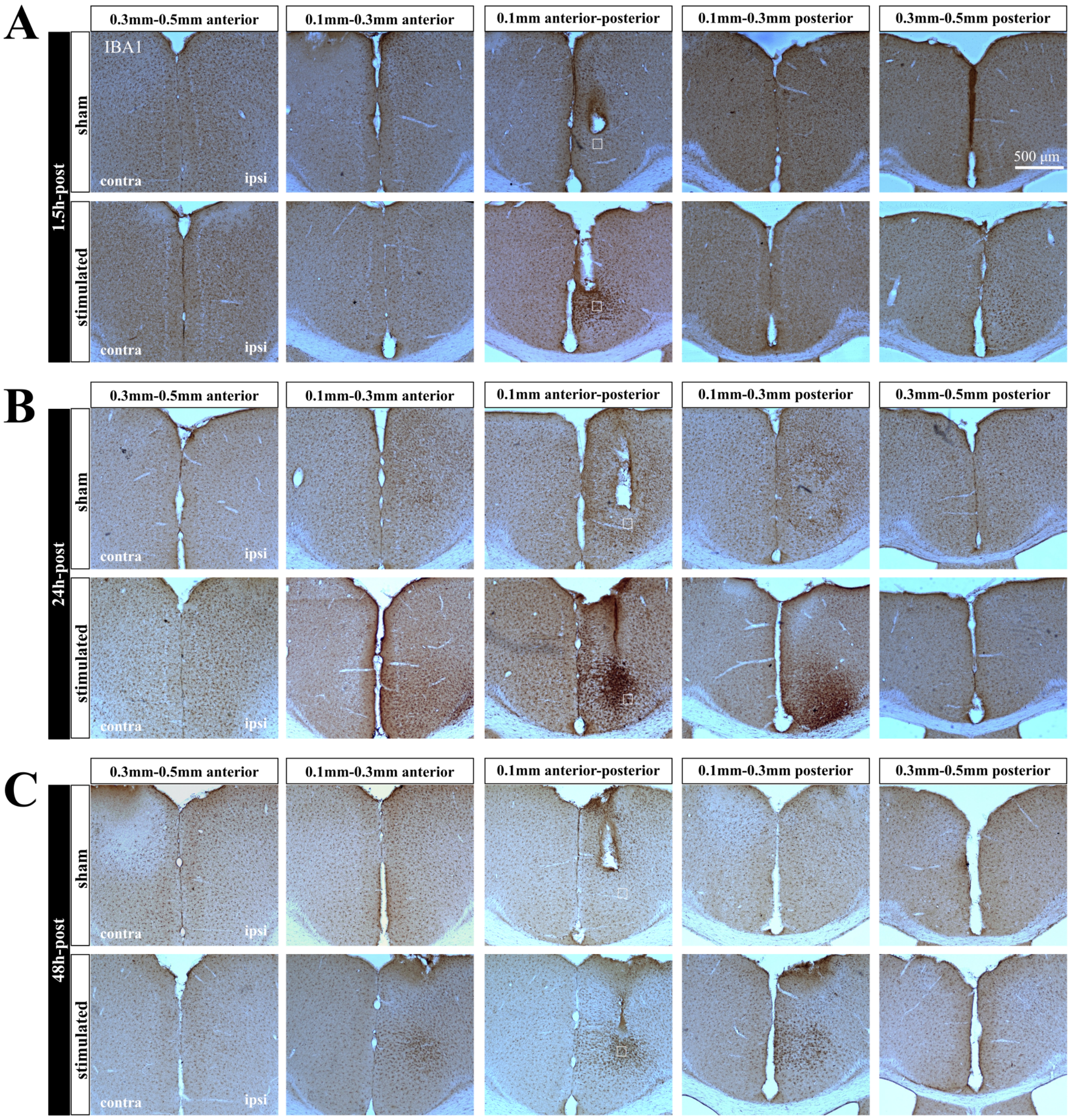


**Supplementary Figure 5-2:** Representative images of IBA1 staining at different distance to the optic fiber at 1.5 h, 24 h, and 48 h post-stimulation. High magnification images of the selected regions (white squares) in the middle panel were presented in Figure 5C.

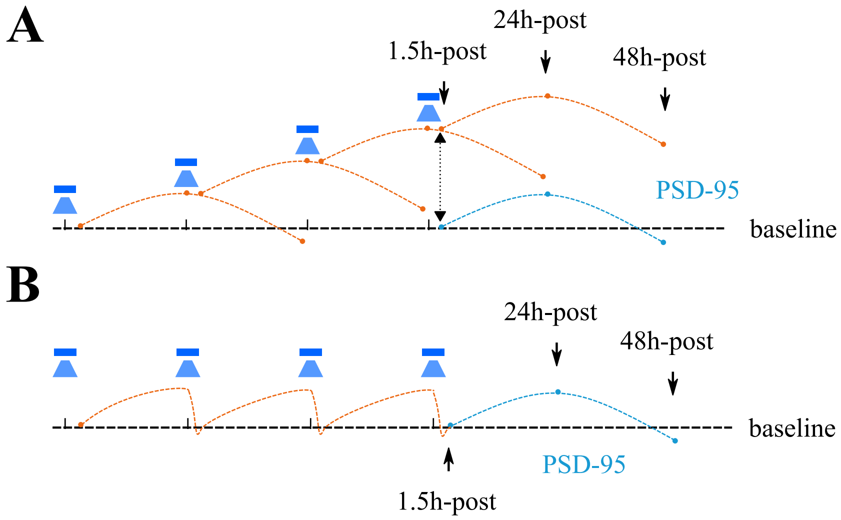


**Supplementary Figure 6-1:** The expected expression of PSD-95 without and with homeostatic regulation. **A** If no homeostatic regulation occurred, PSD-95 expression at 1.5 h after the stimulation should remain upregulated due to accumulated effects. **B** The hypothesized time course of PSD-95 upon each stimulation session.

**Supplementary Table 1:** Animals and experiments used in the current study

| Group | Sample size & genotype | Behavioral tests | Experiments |
|---|---|---|---|
| 1.5 h-post | 12 Thy1-ChR2-YFP (6 shams) | No test | IHC |
| 24 h-post | 19 Thy1-ChR2-YFP (9 shams) | Splash test | IHC |
| 24 h-post | 11 Thy1-ChR2-YFP (5 shams) | Splash test | Microinjection |
| 48 h-post | 10 Thy1-ChR2-YFP (5 shams) | NSF + Splash tests | IHC |
| 48 h-post | 12 Thy1-ChR2-YFP (6 shams) | NSF + Splash tests | Microinjection |
| Virus-injected* | 12 transfected C57BL/6J (6 shams) | NSF + Splash tests | IHC (c-Fos only) |

NSF test, novelty-suppressed feeding test; IHC, immunohistochemical staining.

* Mice were stimulated again after the splash test and sacrificed 1.5 h later.

**Supplementary Table 2:** Primary and secondary antibodies and their concentration used for immunohistochemical staining

| Target Proteins | Primary AB | Primary AB concentration | 2nd AB | 2nd AB concentration |
|---|---|---|---|---|
| VGLUT1 | Millipore, Cat#MAB5502 (RRID: AB_262185) | 1 : 300 | Cy5 donkey anti mouse | 1 : 200 |
| PSD-95 | Millipore, Cat#MAB1596 (RRID: AB_2092365) | 1 : 500 | Cy5 donkey anti mouse | 1 : 200 |
| Neurogranin (Ng) | Millipore, Cat#AB5620 (RRID: AB_91937) | 1 : 1000 | Cy3 donkey anti rabbit | 1 : 200 |
| GFAP | US biological, Cat#2032-27N (RRID: AB_2232293) | 1 : 1000 | Cy3 donkey anti rabbit | 1 : 200 |
| IBA1 | Wako, Cat#019-19741 (RRID: AB_839504) | 1 : 1000 | biotinylated anti rabbit | 1 : 400 |
| c-Fos | Synaptic Systems, Cat#226003 (RRID:AB_2231974) | 1 : 1000 | Cy3 donkey anti rabbit | 1 : 400 |

AB, antibody

**Supplementary Table 3:** Spine classification criteria

| Spine class | Criteria |
|---|---|
| stubby | $\text{length}_{\text{spine}} < 1$ |
| mushroom | $\text{length}_{\text{spine}} < 3$ & $\max(\text{width}_{\text{head}}) > \text{mean}(\text{width}_{\text{neck}}) * 2$ |
| long-thin | $\text{mean}(\text{width}_{\text{head}}) >= \text{mean}(\text{width}_{\text{neck}})$ |
| filopodia | spines that are not stubby, mushroom, long-thin |

**Supplementary Table 4:** Parameters of neuron model

| $\tau_m$ | $t_{\text{ref}}$ | $V_0$ | $V_{\text{reset}}$ | $V_{\text{th}}$ |
|---|---|---|---|---|
| 10.0 ms | 2.0 ms | 0.0 mV | 10.0 mV | 20.0 mV |

**Supplementary Table 5:** Parameters for network model

| $N_E$ | $N_I$ | $\Gamma_{E-I}$ | $\Gamma_{I-E}$ | $\Gamma_{I-I}$ | $J_E$ | $J_I$ | $r_{\text{ext}}$ |
|---|---|---|---|---|---|---|---|
| 10 000 | 2 500 | 10% | 10% | 10% | 0.1 mV | −0.8 mV | 30 kHz |

**Supplementary Table 6:** Parameters for the homeostatic structural plasticity model

| $\epsilon$ | $\nu$ | $\tau_{\text{Ca}}$ | $\beta_{\text{Ca}}$ |
|---|---|---|---|
| 0.008 | $0.004\,\text{s}^{-1}$ | 10 s | 0.0001 |

**Supplementary Table 7:** Parameters used in the model of optogenetic stimulation

| Figure | $r_{\text{opto}}$ (kHz) | $f_{\text{opto}}$ | $T_{\text{opto}}$ (s) | $T_{\text{int}}$ (s) | Repetition | $T_{\text{relax}}$ (s) |
|---|---|---|---|---|---|---|
| 6B | 1.5 | 50% | 75 | - | no | 675 |
| 6E | 1.5 | 50% | 75 | 3 600 | 4 | 7 200 |
